# Development and IND-enabling studies of a novel Cas9 genome-edited autologous CD34^+^ cell therapy to induce fetal hemoglobin for sickle cell disease

**DOI:** 10.1101/2024.04.30.591737

**Authors:** Varun Katta, Kiera O’Keefe, Yichao Li, Thiyagaraj Mayurathan, Cicera R. Lazzarotto, Rachael K. Wood, Rachel M. Levine, Alicia Powers, Kalin Mayberry, Garret Manquen, Yu Yao, Jingjing Zhang, Yoonjeong Jang, Nikitha Nimmagadda, Erin A. Dempsey, GaHyun Lee, Naoya Uchida, Yong Cheng, Frank Fazio, Tim Lockey, Mike Meagher, Akshay Sharma, John F. Tisdale, Sheng Zhou, Jonathan S. Yen, Mitchell J. Weiss, Shengdar Q. Tsai

## Abstract

Sickle cell disease (SCD) is a common severe blood disorder, caused by one major point mutation in the *HBB* gene. Current pharmacotherapies are only partially effective and potentially curative allogeneic hematopoietic stem cell transplantation (HSCT) is associated with immune toxicities. Genome editing of autologous patient hematopoietic stem cells (HSCs) to reactivate fetal hemoglobin (HbF) in erythroid progeny offers a potentially curative approach to treat SCD and circumvents some problems associated with allogeneic HSCT. Although the FDA has released guidelines for evaluating genome editing risks, it remains unclear how to best to assess the preclinical safety and efficacy of genome-edited cellular drug products to prepare for a clinical trial. Here we describe rigorous pre-clinical characterization and optimization of a therapeutic γ-globin gene promoter editing strategy that supported an investigational new drug (IND) application cleared by the FDA. We compared targets in the γ-globin promoter and *BCL11A* erythroid-specific enhancer, identified a lead candidate that potently induces HbF, and tested our approach in mobilized CD34^+^ HSPCs from normal donors and individuals with SCD. We observed efficient editing, induction of HbF to levels predicted to be therapeutic, and reduction of sickling in red blood cells derived from edited HSPCs. With single-cell western and RNA-seq analyses, we defined the heterogeneity and specificity of HbF induction and *HBG1/HBG2* transcription. With CHANGE-seq for sensitive and unbiased genome-wide off-target discovery followed by multiplexed targeted sequencing, we did not detect off-target activity in edited HSPCs. Our study provides a blueprint for translating new discoveries on *ex vivo* genome editing of HSCs towards clinical trials for treating SCD and other blood disorders.

## Introduction

Sickle cell disease (SCD) is a genetic blood disorder most commonly caused by a missense mutation (c.20T>A, p.Glu6Val) in the *HBB* gene that encodes the β-globin subunit of adult hemoglobin (HbA; α2β2).^1^ This mutation causes production of sickle hemoglobin (HbS, α2β^S^2), which polymerizes during hypoxic conditions in venous capillaries, resulting in stiff, brittle sickle-shaped red blood cells (RBCs).^2,3^ SCD is associated with chronic anemia, severe pain crises, progressive multi-organ damage, and early mortality.^4^ Current approved drugs such as hydroxyurea, voxelotor, crizanlizumab and L-glutamine are only partially effective ^5^. Allogeneic hematopoietic stem cell transplantation (HSCT) represents a potential cure but is associated with immune toxicities, including graft-versus-host disease and graft rejection.^6,7^ Moreover, human leukocyte antigen (HLA)-matched donors, which optimize HSCT outcomes, are available for less than 20% of SCD patients. These problems can be circumvented by lentiviral vector-based gene therapy or genome editing of autologous hematopoietic stem cells (HSCs).^8^

Several viral gene therapy and genome editing strategies have been developed to genetically modify autologous HSCs for SCD treatment. These include transduction with lentiviral vectors encoding β- or γ-globin transgenes,^9^ disruption of an erythroid-specific enhancer of *BCL11A,* which represses γ-globin gene transcription,^10^ correction of β-globin gene via homology directed repair (HDR),^11^ and base editing to convert sickle hemoglobin (HbS) to a non-pathogenic variant. ^12^ Two of these strategies have shown promising early results in clinical trials and received FDA approval for use in SCD patients.^9,13^ However, the safest and most effective approach(es) remain to be determined through long-term follow up studies. Each approach has potential genotoxicities, including insertional mutagenesis by lentiviral vectors, unintended off-target activity by genome editing nucleases, bystander edits within the editing window of base editors,^14–16^ genotoxicities associated with base and prime editors, and potential HSC toxicities with high-dose rAAV used to provide DNA template for HDR.^17–19^

Fetal hemoglobin (HbF; α2γ2), the predominant oxygen carrier during late stage fetal development, is a well-established modifier of SCD severity.^20^ Normally around birth, γ-globin expression declines, with a reciprocal increase in the expression of the nearby β-globin gene, leading to a switch from HbF to HbA (α2β2), or HbS in the case of SCD.^21^ Numerous clinical observations have shown that elevated levels of HbF in postnatal RBCs can alleviate SCD pathologies by inhibiting HbS polymerization.^22,23^ Individuals who co-inherit mutations that typically cause SCD along with naturally-occurring mutations that cause hereditary persistence of fetal hemoglobin (HPFH), remain largely asymptomatic.^24,25^

HbF can be induced by using genome editing nucleases such as Cas9 to disrupt repressor-binding sites in the γ-globin promoter.^26,27^ Although Cas9-mediated HDR can correct the mutant sickle *HBB* allele directly, this approach is challenging and may impair HSC engraftment and function.^18^ Moreover, precise edits are accompanied by non-homologous end joining (NHEJ) mediated insertions and deletions (indels) that can disrupt the *HBB* reading frame and reduce the β-globin expression. In contrast, NHEJ-mediated repair can be achieved at high frequencies in quiescent HSCs and can be utilized to re-activate silenced HbF expression.

The perinatal switch from HbF to HbA is mediated by the two repressor proteins BCL11A and ZBTB7A, which bind distinct motifs in the γ-globin promoter to inhibit transcription.^28,29^ Thus, genome editing nuclease-induced indels can be used to re-activate HbF by disrupting an erythroid-specific enhancer in the *BCL11A* gene or by disrupting BCL11A or ZBTB7A binding motifs in the γ-globin gene (*HBG1/HBG2*) promoters.^27,30^ A clinical trial examining Cas9-mediated disruption of the *BCL11A* enhancer shows promising early results in patients with SCD or β-thalassemia.^10,31,32^

Previously, we showed that Cas9 disruption of a BCL11A repressor-binding site (113 to 118 nucleotides upstream of the transcription start site in the γ-globin gene promoters) in HSCs resulted in the induction of HbF to potentially therapeutic levels in RBC progeny generated in *in vitro* and *in vivo*.^27,33^ Potential advantages of our approach include transient Cas9 expression for *ex vivo* modification of autologous HSCs and high efficiency of NHEJ-mediated γ-globin promoter editing compared to HDR correction of the HbS mutation. Moreover, disruption of the BCL11A binding site recapitulates benign HPFH mutations and may have more specific biological effects compared to the global loss of *BCL11A* expression in RBCs.^34,35^

Here we show a set of pre-clinical experiments characterizing and advancing our genome editing approach towards a clinical trial. Cas9 disruption of the −115 BCL11A binding motif in the γ-globin promoter, hereafter referred to as −115 γ-globin promoter, induced HbF more potently than Cas9 disruption of the upstream ZBTB7A binding motif. Similarly, disruption of BCL11A motif in SCD donor CD34^+^ hematopoietic stem progenitor cells (HSPCs) achieved consistent, high-frequency editing of repopulating HSCs and substantial induction of red blood cell HbF. Single-cell analysis by western blotting and RNA-seq showed that HbF was induced in most donor-derived erythroid cells with significant reductions in sickling after exposure to hypoxia. No off-target indels or genomic re-arrangements were detected using state-of-the-art, sensitive, unbiased genome-wide assays and targeted sequencing. Overall, our preclinical studies suggest that Cas9-mediated disruption of the BCL11A repressor binding site in the γ-globin promoters represent a promising approach for genomic therapy of SCD and β-thalassemia.

## Results

### Disrupting the γ-globin promoter *(HBG1/HBG2*) BCL11A-binding motif efficiently induces HbF

We compared the magnitude of HbF induction after Cas9 nuclease disruption of the BCL11A and ZBTB7A motifs in the γ-globin promoter which each harbor naturally occurring HPFH mutations. ^36^ We electroporated peripheral-blood mobilized CD34^+^ hematopoietic stem progenitor cells (HSPCs) from normal donors with Cas9 ribonucleoprotein complexes (RNPs) targeting the ZBTB7A or BCL11A binding motifs to induce DNA double strand breaks (DSBs) at positions −196 or −115, respectively (**Figure 1A**). We tested both wild-type Cas9 and a high-fidelity version, ^37^ each containing an enhanced nuclear localization signal that improves editing efficiency (SpCas9-3xNLS).^38^ Four days after electroporation (EP) of wild-type Cas9-3xNLS RNP, measured average indel frequencies were 76.2% and 85.7% at the ZBTB7A and BCL11A motifs, respectively. Editing with high-fidelity Cas9-3xNLS RNP resulted in 79.7% and 83.3% of indels at the ZBTB7A and BCL11A motifs, respectively (**Figure 1B**). Thus, under protein-specific optimized transfection conditions (see methods), wild-type and high-fidelity Cas9-3xNLS generated similar indel frequencies at the two target sites in CD34^+^ HSPCs.

**Figure 1.**
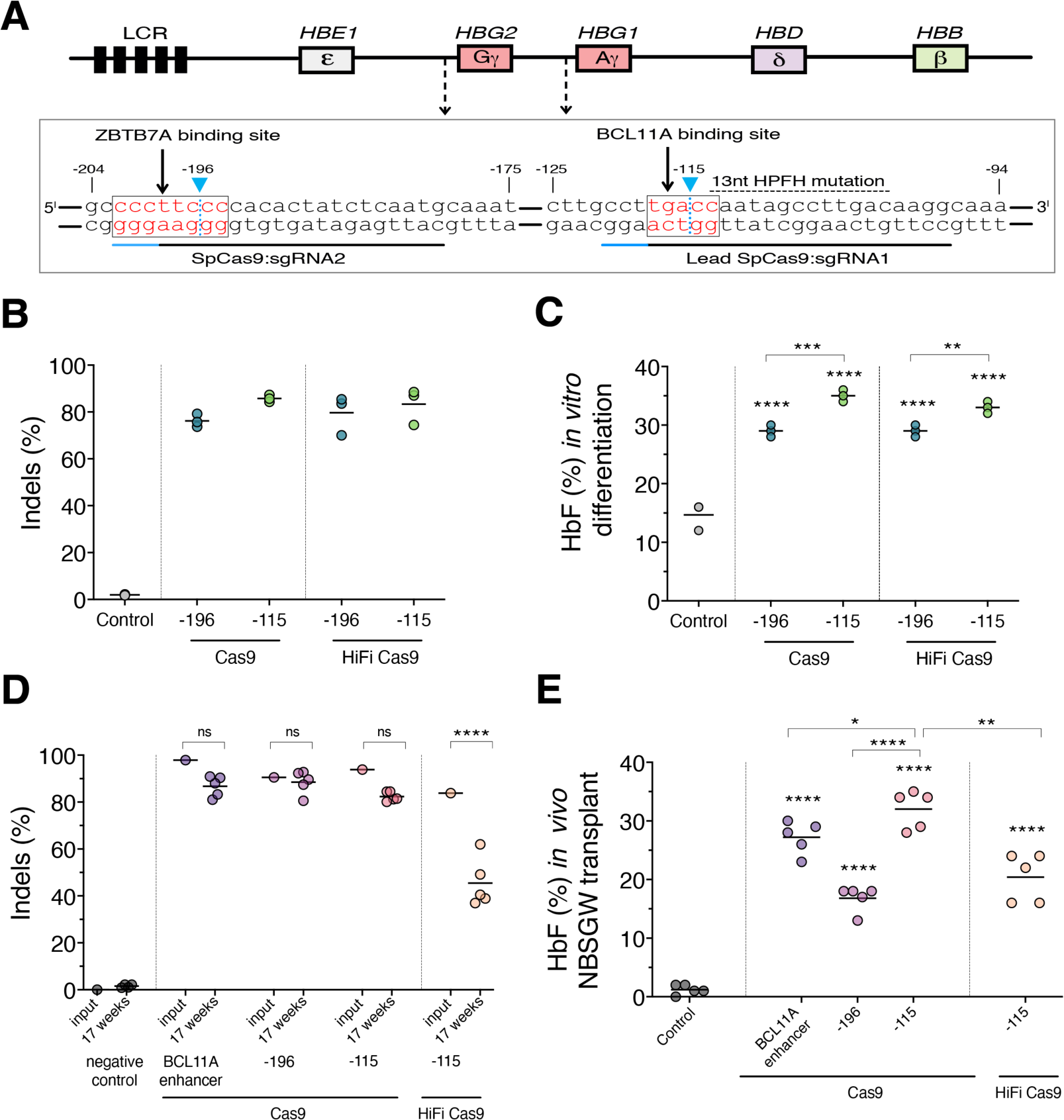
HbF is more effectively induced by editing of −115 BCL11A-binding motif than −196 ZBTB7A binding motif in the γ-globin promoters. (**A**) Schematic of SpCas9-3xNLS genome editing target sites in the γ-globin promoters highlighting ZBTB7A (−196) and BCL11A (−115) binding motifs in red (boxed). Blue arrow with dotted line shows the predicted SpCas9 cleavage position at −196 and −115 sites. Single guide-RNA design location shown in black line with SpCas9 protospacer adjacent motif (PAM) highlighted in blue line (NGG) for respective sites. 13nt deletion associated with hereditary persistence of fetal hemoglobin (HPFH) mutation indicated with black dotted line. (**B**) Indels for WT SpCas9 and HiFi SpCas9 (R691A) determined by NGS after editing at −196 and −115 γ-globin promoter targets using Lonza 4D nucleofector under optimal conditions. Scatter plots shown as mean with error line. (**C**) Percent of HbF estimated in erythroid differentiated cultures at Day 21 by ion exchange high-performance liquid chromatography (HPLC). Scatter plots show the results as a mean, adjusted P values for all samples are <0.0001**** compared to controls (ordinary one-way ANOVA, Dunnett’s multiple comparison tests). Adjusted P values for −196 vs −115 γ-targets with WT and HF are 0.0002*** and 0.0024** (2way ANOVA, Šídák’s multiple comparisons test). (**D**) On-target indels determined by NGS after editing at day 4 and 17 weeks of post-transplantation for different targets *BCL11A* enhancer, −196 γ-globin promoter, and −115 γ-globin promoter (‘ns’ represents statistically not significant for input and 17 weeks, P values 0.374, 0.991, 0.347 and −115 HF γ-globin promoter (P value <0.0001****) (2way ANOVA, Šídák’s multiple comparisons test). Each dot represents an independent mouse from 17 weeks (n = 5). (**E**) Scatter plot of %HbF in human erythroid cells for control or edited CD34^+^ cells, measured by HPLC at 17 weeks of xenotransplantation. Data shown as mean with error line and adjusted P value <0.0001**** (n=5) compared to controls as each value shown as dots (Ordinary one-way ANOVA, Dunnett’s multiple comparison tests) and adjusted P values for *BCL11A* enhancer vs −115 with WT, −196 vs −115 with WT, and −115 with WT vs HF are 0.0169*, <0.0001****, and 0.0001*** (Ordinary one-way ANOVA, Tukey’s multiple comparison tests).

To quantify HbF induction after editing, we generated late-stage erythroblasts by *in vitro* differentiation, followed by high performance liquid chromatography (HPLC) of cell lysates. Editing at either γ-globin promoter target resulted in significantly increased HbF compared to unedited controls (**Figure 1C**). Induction of HbF was significantly higher after wild-type Cas9-3xNLS editing of the −115 γ-globin promoter 35% (1.00), mean (SD) compared to editing at the −196 γ-globin promoter 29% (1.00) (adjusted P value 0.0002). We obtained similar results with high-fidelity Cas9-3xNLS *in vitro* (**Figure 1C**).

To determine editing outcomes in bone marrow (BM) repopulating HSCs, which represent a small proportion of human CD34^+^ cells, we transplanted normal donor CD34^+^ HSPCs edited with wild-type Cas9 at the −196 and −115 γ-globin promoter targets, and CD34^+^ HSPCs edited with high-fidelity Cas9 at −115 γ-globin promoter target into immunodeficient NBSGW mice. We also analyzed CD34^+^ HSPCs edited at the erythroid-specific enhancer of *BCL11A* as a positive control, and unedited CD34^+^ HSPCs as a negative control. At 17 weeks post-transplantation, we collected human donor cells from the BM of transplanted mice and measured engraftment, multi-lineage differentiation, indel frequencies, and erythroid HbF expression. We analyzed erythroblasts and reticulocytes isolated from the BM of transplanted mice because human RBCs have a very short half-life in mouse circulation.^39^ Indels in input CD34^+^ HSPCs prior to transplantation were 83.8% to 97.9% at the *BCL11A* enhancer, −115, and −196 γ-globin promoter targets (**Figure 1D**).

After transplantation, repopulating human HSPCs edited with wild-type Cas9 retained high indel frequencies: 86.7% (4.37), 88.5% (4.95), and 82.4% (2.00) at the *BCL11A* enhancer, −196, and −115 γ-globin promoter targets respectively. In contrast, indels were significantly reduced in repopulating CD34^+^ HSPCs edited at the −115 γ-globin promoter target with high-fidelity Cas9 45.5% (10.3) vs input 83.8%, (adjusted P value <0.0001) compared to those edited with wild-type Cas9 (**Figure 1D**). Despite similar indels with wild-type and high-fidelity Cas9 in input cells prior to transplantation, some indel mutations (such as −1, −2, −3, −4) associated with high-fidelity Cas9 declined after 17 weeks (**Figure S1A and S1B**). We conclude that wild-type Cas9 more efficiently edits repopulating HSCs than high-fidelity Cas9 at our −115 γ-globin promoter target. These studies indicate that wild-type Cas9 more efficiently edits repopulating HSCs than high-fidelity Cas9 at our −115 γ-globin promoter target.

To compare HbF induction after editing the −115 and −196 γ-globin promoter target sites, we analyzed CD235a^+^ erythroid progeny of transplanted CD34^+^ HSPCs by HPLC. HbF induction with wild-type Cas9 at the −115 γ-globin promoter target 32% (3.24) was significantly higher compared to unedited controls (<2%, adjusted P value <0.0001). In contrast, editing at the −196 γ-globin promoter target resulted in significantly less HbF induction 16.8% (2.17), adjusted P value <0.0001) compared to the −115 γ-globin promoter target. As expected, lower editing observed with high-fidelity Cas9 at the −115 γ-globin promoter target resulted in corresponding lower HbF induction 20.4% (4.1) (adjusted P value 0.0001) compared to wild-type Cas9. Moreover, −115 γ-globin promoter editing induced higher levels of HbF 32% compared to the cells edited at the *BCL11A* enhancer 27.2% (2.78) (adjusted P value 0.0169) (**Figure 1E**). These results suggest that −115 γ-globin promoter editing could be an effective alternate strategy for HbF induction with higher biological specificity.

Taken together, we conclude that editing at the −115 γ-globin promoter target with wild-type Cas9 represents the most potent nuclease-based approach for therapeutic HbF induction and chose this strategy for further evaluation as a therapeutic lead candidate.

### Therapeutically relevant levels of HbF with reduction of RBC sickling

To assess the utility of our genome editing approach for SCD, we electroporated plerixafor-mobilized CD34^+^ HSPCs from three individuals with SCD (2 with HbSS and 1 with HbSC genotype) and 1 normal donor with Cas9-3xNLS RNP targeting the −115 γ-globin promoter, followed by transplantation into NBSGW mice. After 17 weeks, human cell engraftment in mouse BM was 90.6% (3.32), as measured by expression of the human CD45^+^ hematopoietic antigen. Engraftment levels of unedited control HSPCs were similar 90.9% (3.65) (**Figure 2A**). We observed no significant differences in the differentiation of edited or control HSCs to T-cell, B-cell, myeloid or erythroid lineages in vivo (**Figure 2B**). Human donor cell erythroid maturation measured by the expression of stage-specific markers CD49d, CD235a, Band3 and Hoescht staining to identify reticulocytes, was similar in edited and control samples (**Figure S2A and S2B**).

**Figure 2.**
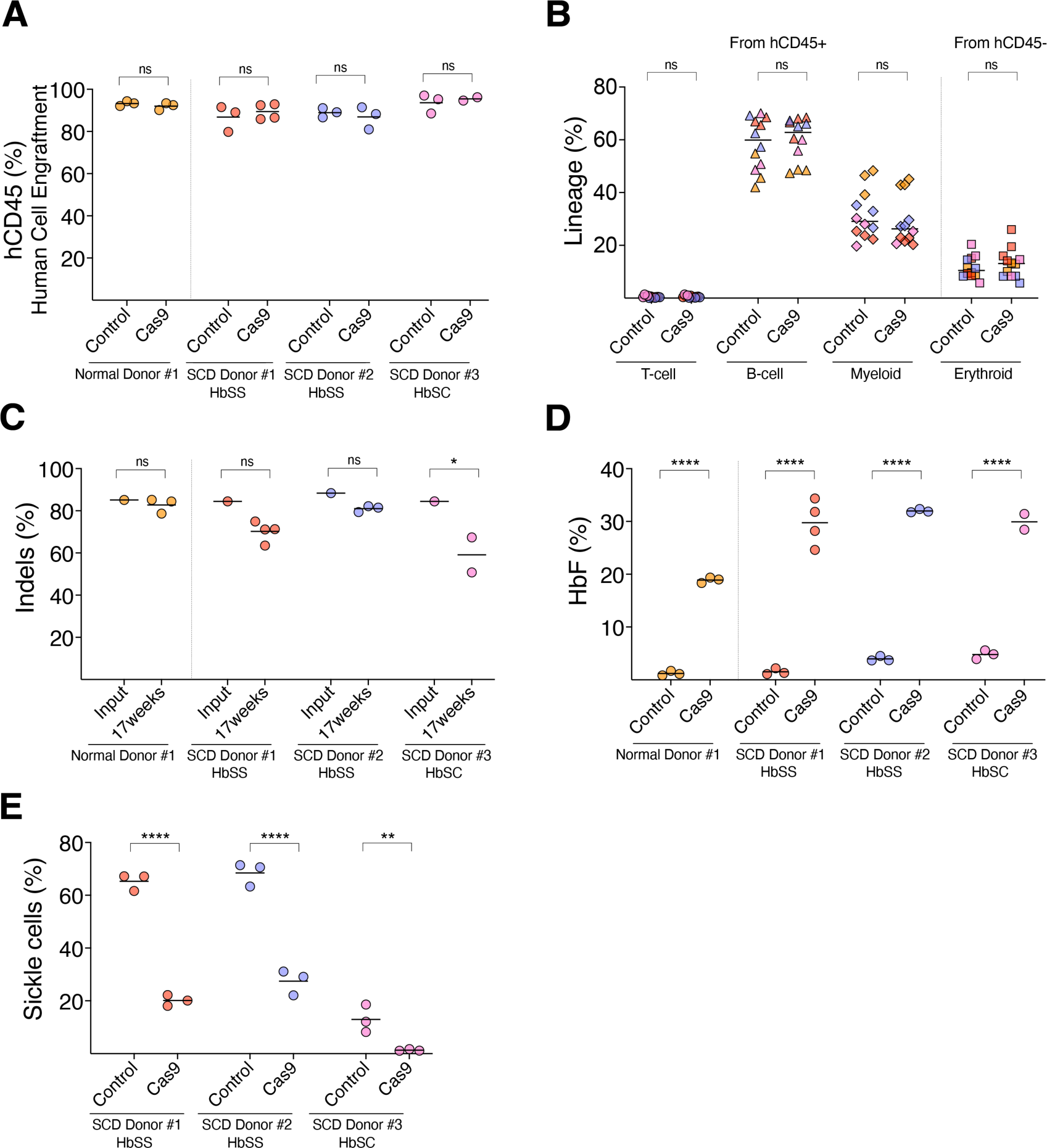
Editing of −115 γ-gamma globin promoter efficiently induces fetal hemoglobin and functionally reduces sickling in erythroid progeny of edited CD34+ HSPCs from SCD patients. Plerixafor-mobilized CD34^+^HSPCs derived from normal donor and 3 SCD donors electroporated with WT-Cas9-3xNLS and unedited controls were transplanted into mice and analyzed 17 weeks later. Each donor is annotated with different color (orange, red, purple, pink) and each dot in the graph represents an individual mouse with average indicated by solid line. (**A**) Percentage of human engraftment (hCD45^+^) evaluated in mouse BM after 17 weeks by flow cytometry (P values -−0.992, 0.855, 0.948, and 0.977 and ‘ns’ indicates statistically insignificant, 2way ANOVA, Šídák’s multiple comparisons test). (**B**) Percentages of human donor-derived lineages shown as human T (CD3^+^), B (CD19^+^), and Myeloid (CD33^+^) cells in human CD45^+^ population in mouse BM at 17 weeks and erythroid lineage is shown as the %CD235a^+^ cells within the CD45^−^ population from mouse and human. P values are >0.999, 0.959, 0.862, 0.9755 and statistically not significant compared to controls (2way ANOVA, Šídák’s multiple comparisons test). (**C**) On-target indels measured by high-throughput sequencing 4 days after editing (infusion product/input) or after xenotransplantation into mice at 17 weeks (P values for normal, HbSS #1, and HbSS #2 are 0.993, 0.177, 0.723, not significant and SCD donor #3 is 0.021* (2way ANOVA, Šídák’s multiple comparisons test). (**D**) Percentages of HbF in human erythroid cells isolated from mouse BM measured by HPLC. Adjusted P values are <0.0001**** and significant (2way ANOVA, Šídák’s multiple comparisons test). (**E**) In vitro differentiated human erythroid cells derived from edited, and controls were sorted for enucleated cells and incubated for 8 hours in 2% O_2._ Enumerated % sickled reticulocytes from the images shown in **fig. S3A** and more than 300 cells/image from three technical replicates counted by 2 blinded observers. P < 0.0001****, P < 0.0001****, and P < 0.0075** for HbSS, HbSS, and HbSC donors (2way ANOVA, Šídák’s multiple comparisons test).

To determine −115 γ-globin promoter editing frequencies before and after NBSGW mouse transplant, and associated levels of HbF induction, we performed targeted next-generation sequencing (NGS) and analyzed HbF levels in CD235a^+^ erythroid cells by HPLC. We achieved consistent high-frequency editing 85.6% (1.87) in CD34^+^ HSPCs in pre-transplantation input cells from one normal and three SCD donors. Seventeen weeks after transplant, indels remained were 70.1% (10.96) in repopulating human cells from SCD donors (**Figure 2C**), which resulted in 30.6% (1.24) HbF induction in erythroid progeny isolated from mouse BM (**Figure 2D**).

To assess the functional effects of HbF induction in RBCs, we edited the −115 γ-globin promoter target in CD34^+^ cells from SCD donors, generated reticulocytes by *in vitro* erythroid differentiation and exposed them to low oxygen concentration (2%) for 8 hours. Phase contrast microscopy showed that reticulocytes derived from unedited HSPCs exhibited 66.9% (2.22) sickled morphology, whereas sickling was significantly reduced to 23.8% (5.16) in reticulocytes derived from edited cells from two donors with HbSS genotype. In reticulocytes derived from a HbSC donor HSPCs, the fraction of sickled reticulocytes was 12.9% (5.25) in unedited controls and 1.3% (0.34) after editing (**Figures 2E and S3A**). These results demonstrate that disruption of the −115 γ-globin promoter target induces HbF expression to levels that ameliorate sickling *in vitr*o, therefore predicting therapeutic efficacy.

### Broad induction of HbF in red blood cell progeny

Optimal therapeutic induction of γ-globin requires that most or all RBCs express HbF above a threshold level that inhibits HbS polymerization.^40,41^ Thus, ideal genome editing therapies for SCD should induce both high and broad (i.e., pancellular) expression of HbF.^42^

We used three orthogonal methods to evaluate HbF expression in erythroid cells generated from edited or control SCD donor HSPCs after xenotransplantation. First, we used flow cytometry to measure HbF immunostaining erythroid cells (F-cells). We observed that 74.4% (3.84) of erythroblasts expressed HbF, albeit with relatively high background in control cells 29.8% (2.26) (**Figure 3A**) which is commonly observed in this assay.^33^ Second, we performed single cell western (scWestern) analysis. Pooled CD235a^+^ erythroid cells from each donor were immobilized on a polyacrylamide gel-chip, lysed, fractionated by sodium dodecyl-sulfate polyacrylamide gel electrophoresis (SDS-PAGE) and probed with anti-γ-globin and anti-α-globin antibodies. In contrast to flow cytometry, scWestern blot analysis detected γ-globin expression in 49.3 % to 58.2% of edited erythroid cells, whereas <6.1% of unedited control cells expressed HbF (**Figure 3B**). While the immune-flow cytometry and scWestern analysis detected similar relative changes in HbF-expressing cells after editing, the absolute numbers of F-cells detected by scWestern was slightly lower than the numbers detected by flow cytometry. This apparent discrepancy may reflect differences in the relative sensitivity and dynamic range of the two techniques.

**Figure 3.**
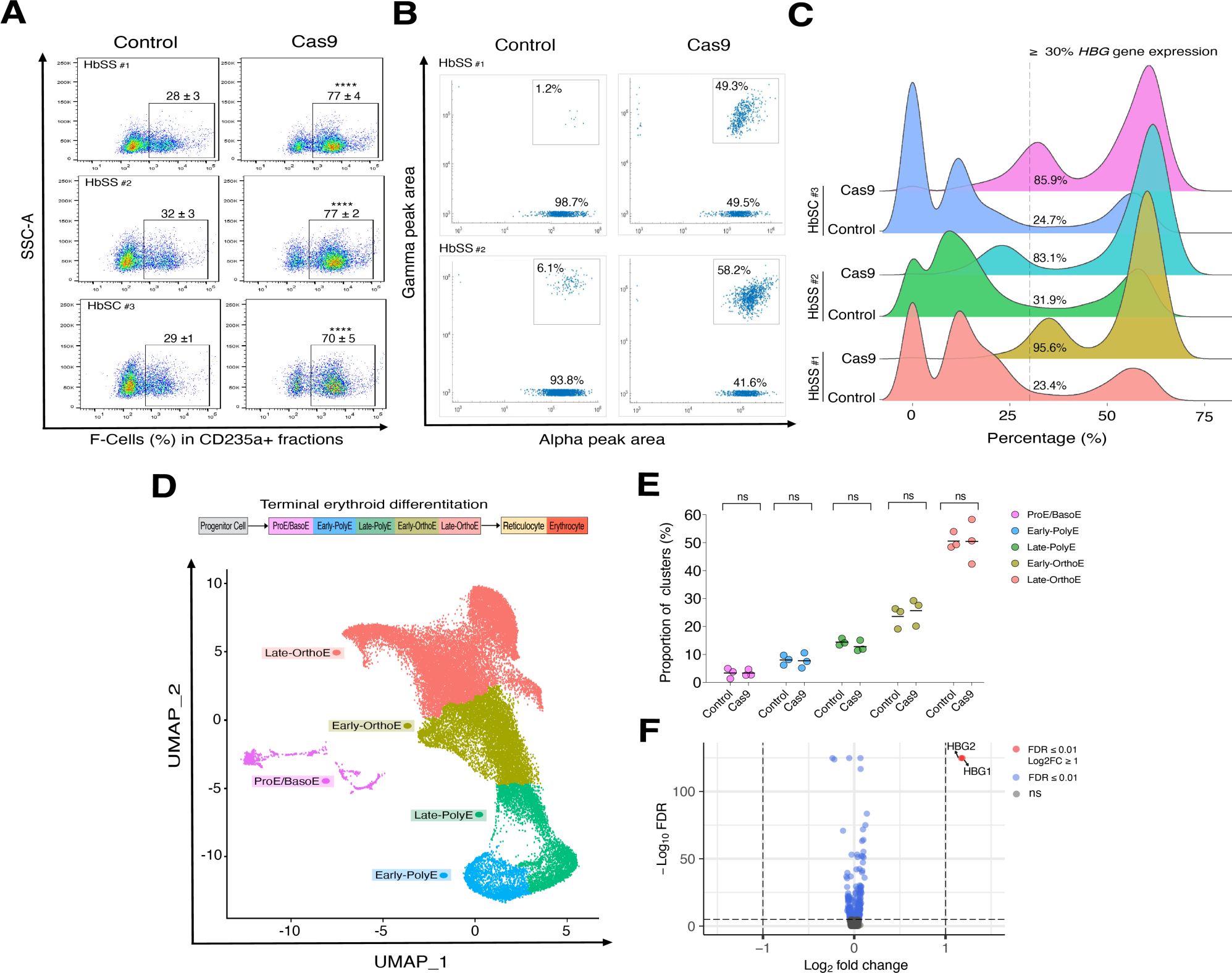
Editing 115 γ-globin promoter site broadly induces fetal hemoglobin as measured by complementary single-cell analyses. Analysis of CD235a^+^ cells from transplanted mouse BM derived from three SCD donors at 17 weeks. (**A**) Representative scatter plots of F-expressing cells by flow cytometry for control and Cas9 edited cells. Percentages were shown as mean (SD), with P values are <0.0001**** and significant for all three donors (2way ANOVA, Šídák’s multiple comparisons test). (**B**) Scatter plots of α-globin (shown on x-axis) and γ-globin (shown on x-axis) comparing between control and Cas9-3xNLS by single cell western analysis. (**C**) Kernel density plots showing the distribution of the percentage of *HBG transcripts* (*HBG1*+*HBG2*/*HBG1*+*HBG2*+*HBB*) in Cas9 edited erythroblasts compared to unedited controls. (**D**) UMAP plot showing annotated cell clusters at different stages of erythroid maturation from pooled erythroid samples derived from mice BM. Terminal erythroid differentiation stages classified as proerythroblast/basophilic erythroblast (ProE/BasoE), early & late stages of polychromatophilic erythroblast (PolyE), and early & late stages of orthochromatic erythroblast (OrthoE). (**E**) Scatter plot showing the percentage of each cluster at different stages of erythroid differentiation between control and Cas9. Differences between control and Cas9 were not significant (i.e., ns) (2way ANOVA, Šídák’s multiple comparisons test). (**F**) Volcano plot showing differentially expressed genes. Blue dot indicates the expressed genes with FDR ≤ 0.01, red dot indicates upregulated genes with both FDR ≤ 0.01 and log_2_FC ≥ 1, and grey dot (ns) indicates not significant.

To better understand the differences in detection of HbF protein in the two single cell assays, we analyzed the relative levels of γ-globin transcript by single cell RNA sequencing (scRNA-seq). Transcription of the γ-globin genes correlates with HbF protein synthesis in erythroid cells.^43^ γ-globin mRNA was increased in edited erythroid cells, with an average of 88.2% (6.62) expressing more than 30% γ-globin mRNAs relative to all β-like globin transcripts including *HBB* (β-globin) and *HBD* (δ-globin). Consistent with flow cytometry analysis, 26.7% (4.59) of unedited erythroid cells derived from SCD donor HSPCs expressed more than 30% γ-globin mRNA. Interestingly, we observed a bimodal distribution of *HBG* expression that was clearly shifted in edited cells compared to unedited ones (**Figure 3C**). Our results suggest that most edits induce γ-globin transcription into a therapeutically relevant range, but some more potently than others.

To determine whether editing altered the distribution of erythroblast subpopulations, we integrated scRNA-seq data from all cells and performed dimensionality reduction using uniform manifold approximation and projection (UMAP)^44^ (**Figure 3D**). We clustered the cells according to standard erythroid maturation stages including proerythroblast (ProE), basophilic erythroblast (BasoE), polychromatophilic erythroblast (PolyE), orthochromatic erythroblast (OrthoE) and reticulocyte based on gene specific markers^45^ (**Figure S4A**). Editing did not significantly alter the relative proportions of cells at these maturation stages (**Figure 3E**).

To further determine whether editing of HSCs altered transcription in erythroid progeny we performed differential expression analysis to compare edited and control cell populations. *HBG1* and *HBG2* mRNA levels were significantly and specifically upregulated in edited cells (**Figure 3F, and S5A**). No differences in other erythroid-expressed mRNAs including *SLCA1, ITG4,* and *GYPA* were detected in edited vs unedited cells. (**Figure S6A**).

We note that the levels of HbF indicated by our analysis of human SCD donor-derived erythroid cells in mouse BM likely underestimate the levels that would be expressed human subjects who receive autologous edited HSCs because high levels of HbF reduce HbS polymerization and increase the lifespan of circulating RBCs.^46^ This positive selection of HbF-expressing RBCs cannot be measured in human-to-mouse xenotransplantation studies because human RBCs are rapidly eliminated from the mouse circulation.^39^

Overall, single cell analysis of γ-globin protein and mRNA shows that the −115 γ-globin promoter edit induces HbF expression broadly and to levels that are predicted to offer therapeutic benefit.

### Good manufacturing practice (GMP) production and characterization of critical reagents: Cas9, gRNA, Cas9 RNP complex

To manufacture GMP-grade Cas9-3xNLS, we developed a purification protocol without affinity tags, which are undesired for therapeutic protein production. *E.coli* T7 express cells containing Cas9 expression plasmid were grown in a 5L fermenter, and Cas9 expression was induced for 22 hours. Cell pellets from the fermenter were homogenized and processed to purify high-quality clinical-grade Cas9 protein. Downstream steps included Sepharose SP column purification, allantoin addition for endotoxin removal, and ceramic hydroxyapatite type I column purification (**Figure 4A**). The resulting Cas9 protein passed release testing with acceptable levels of residual *E. coli* host cell protein (3.47ng/mg protein) and endotoxin (0.34 EU/mg protein). The purity of Cas9 protein was >99% as evaluated by Reverse-Phase Ultra-high-Performance Chromatography (RP-UPLC), Sodium Dodecyl-Sulfate Capillary Gel electrophoresis (SDS-CGE), and Size Exclusion Chromatography (SEC) (**Figure 4B**).

**Figure 4.**
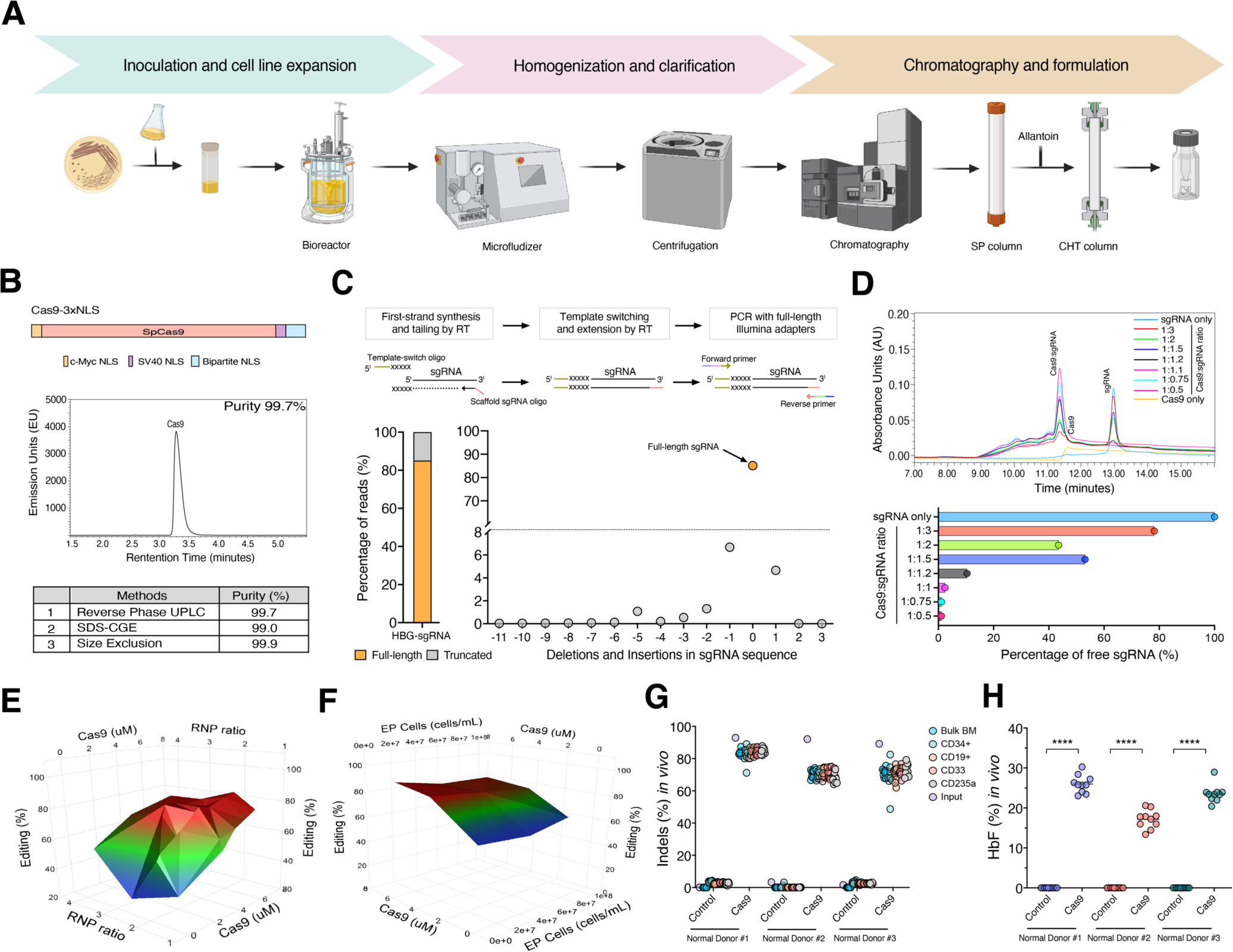
Development of a cGMP, clinical-scale process for −115 γ-globin promoter editing. (**A**) Streamline for Cas9-3xNLS purification under GMP conditions. (**B**) Schematic version of Cas9 protein with c-Myc nuclear localization signal (NLS) at N-terminal and both SV40 NLS and bipartite NLS tags at C-terminal. Reverse Phase UPLC analysis of purified Cas9-3xNLS fraction and purity of Cas9 evaluated by different methods. (**C**) Illustration of single guide-RNA (sgRNA) sequencing of GMP-grade HBG-sgRNA using smarter smRNA-seq technology. Bar and scatter plots show the percentage of reads mapped to full-length targeted sgRNA (shown on y-axis) and insertions and deletions in the sgRNA sequence (shown on x-axis). (**D**) Size exclusion chromatography for RNP complexes and free sgRNA residues at different molar ratios of Cas9:sgRNA ranging from 1:0.5 to 1:3. Bar plot represents the percentages of free sgRNA at different Cas9-3xNLS:sgRNA ratios. Surface plot of optimization conditions through DOE studies using JMP, looking at editing efficiency with (**E**) Cas9 concentration and RNP ratio, and (**F**) Cas9 concentration and cell concentration. (**G**) Measurement of indels in all hematopoietic lineages by NGS analysis before transplant (input) and after transplant in bulk BM, CD34^+^, CD33^+^ (myeloid), CD19^+^ (B-cell) and CD235a^+^ (erythroid) using MaxCyte GTX (n=10). (**H**) Ion exchange-HPLC analysis for HbF induction in erythroid cells after 17 weeks post transplantation (n=10). Adjusted P values are <0.0001**** for control vs cas9 (2way ANOVA, Šídák’s multiple comparisons test).

The single guide RNA (sgRNA) includes two components: crispr RNA (crRNA) a 20bp complementary sequence that binds to the targeted region and an 80bp trans-activating crRNA (tracrRNA) that serves as a scaffold for Cas9 ribonucleoprotein complex formation.^47^ Commercially synthesized sgRNAs used in this study including GMP-grade HBG-sgRNA are chemically modified to increase the stability and editing efficacy.^48^ The quality of sgRNA is critical for therapeutic genome editing, as heterogeneous contaminant RNAs could impair on-target editing or increase off-target edits at cognate target sites. We utilized a modified SMART smRNA sequencing protocol to analyze the −115 γ-globin targeting sgRNA.^49^ 85.1% of the sequencing reads generated from commercially synthesized, GMP-grade sgRNA aligned perfectly to the −115 γ-globin promoter target sequence, while the remaining 14.9% contained truncations in targeting and/or scaffold regions that could impact the efficacy or specificity of editing (**Figure 4C**). From the mapped reads, 80.79% of the 5’ protospacer sequence perfectly matched the targeting sgRNA sequence with complementarity to the *HBG* promoter and 64.9% matched the full-length product (**Figure S7A**). Most importantly, we did not detect evidence of contaminating gRNAs that could target other regions of the genome, consistent with full GMP changeover procedures (**Figure S7B**).

To optimize Cas9 RNP complexation and minimize the amount of free sgRNA that could induce an innate immune response, we titrated the molar ratios of Cas9 to sgRNA. Using the same buffers and protocol as in GMP manufacturing runs, we generated RNP complexes with different Cas9:sgRNA ratios followed by size exclusion chromatography with HPLC-UV spectroscopy to quantify the levels of RNP complex, Cas9 and free sgRNA. A Cas9:sgRNA ratio of 1:1.2 resulted in complexation of most Cas9 with 10.4% free sgRNA, whereas increasing Cas9:sgRNA ratios from 1:1.5, to 1:3 ratios resulted in 43.5% to 78.2% free sgRNA (**Figure 4D**). This provided a Cas9 complexation range for further cell-based optimization of editing efficiency and cell recovery.

To develop an efficient clinical-scale process for editing CD34^+^ HSPCs at −115 γ-globin promoter, first, we compared EP programs for transfecting RNP using the MaxCyte GTX. HSC3 and HSC4 programs resulted in comparable editing frequencies and cell viabilities (**Figures S8A and S8B)**. Thus, we selected program HSC3 for EP, which uses a lower energy program and may reduce electroporation-associated cell toxicity. Next, we performed design-of-experiments (DOE) to optimize Cas9 concentration, Cas9:sgRNA complexation ratio, and cell concentration during EP. Editing efficiency *in vitro* plateaued at 4.8 μM Cas9 protein and a Cas9:sgRNA ratio of 1:1.2 (**Figure 4E**). Editing efficiencies and cell survival were similar at cell densities of 10-100 million/ml during EP (**Figure 4F**). Additionally, we determined that the editing efficiency was highest after pre-stimulating plerixafor-mobilized CD34^+^ HSPCs for one day prior to EP (**Figure S8C**).

In principle, residual Cas9 protein in transplanted HPSCs could stimulate a host immune response.^50^ We performed western blot analysis and measurement of on-target indels across multiple timepoints to assess the kinetics of Cas9 expression and genome editing. After electroporation of RNP into HSPCs, Cas9 activity measured by indel formation reached a plateau by day 4 and no Cas9 protein was detected after 7 days (**Figure S8D**). However, in CD34^+^ HSPCs cryopreserved after 24 hours of EP, Cas9 was completely undetectable 72 hours in post-thaw edited cells (**Figure S8E**). These results show that intracellular Cas9 is transient and fully degraded by day 3.

We combined optimized EP parameters and cell purification process to conduct a large scale pre-GMP engineering run with plerixafor-mobilized normal donor CD34^+^ HSPCs (n=3). For this scale-up, we used GMP-like sgRNA (Synthego) and GMP-like Cas9 protein manufactured from our GMP facility. We conducted three pre-GMP engineering runs and transplanted edited and unedited control cells into NBSGW mice for in vivo pharmacology studies. At 17 weeks post-transplantation, the indels in donor-derived BM, CD34^+^, CD19^+^, CD33^+^ and CD235a^+^ cells were similar to each other and to that of the input CD34^+^ cells measured 5 days after editing (**Figure 4G**). In donor-derived CD235a^+^ erythroid cells HbF levels were 17.2% to 26% after editing compared to 0% in unedited control cells (**Figure 4H**). The fraction of F-cells was 55.1% to 73.3% after editing compared to 11.9% to 20.2% in unedited cells (**Figure S8F**). As obvious, there was no change in enucleation between the percentages of edited and unedited control cells (**Figure S8G**).

Altogether, we establish a framework for clinical scale-up process to manufacture and characterize critical genome editing reagents like Cas9 and sgRNA, and an edited CD34^+^ cellular drug product. Next, we achieve efficient editing in normal CD34^+^ HSPCs at a large scale to induce similar and effective levels of HbF to further support the efficacy of our preclinical studies.

### Assessment of genetic outcomes of Cas9-mediated disruption of −115 γ-globin promoters

Recent draft U.S. Food and Drug Administration (FDA) guidelines ^51^ highlighted potential safety concerns specifically associated with genome editing. Genome-wide genotoxicities can include small off-target indels and structural rearrangements such as large deletions, inversions, and translocations.

Simultaneous on-target Cas9 indels in the tandem, ancestrally duplicated *HBG1 and HBG2* genes can result in deletion of the intervening ∼4.9 kb region to generate a single hybrid gene containing the upstream *HBG2* promoter sequence fused to the downstream *HBG1* promoter and coding sequences (**Figure 5A**). To detect these large 4.9 kb deletions in the β-globin gene cluster locus, we used commercially available Multiplex Ligation Dependent Probe Amplification (MLPA) assay. Five probes that bind to the aligned map locations 11-005 227321, 11-005 227546, 11-005 228965, 11-005 230885, 11-005 226171 have shown possible copy number variations (CNVs) in bulk edited BM transplants (**Figure S9A**).

**Figure 5.**
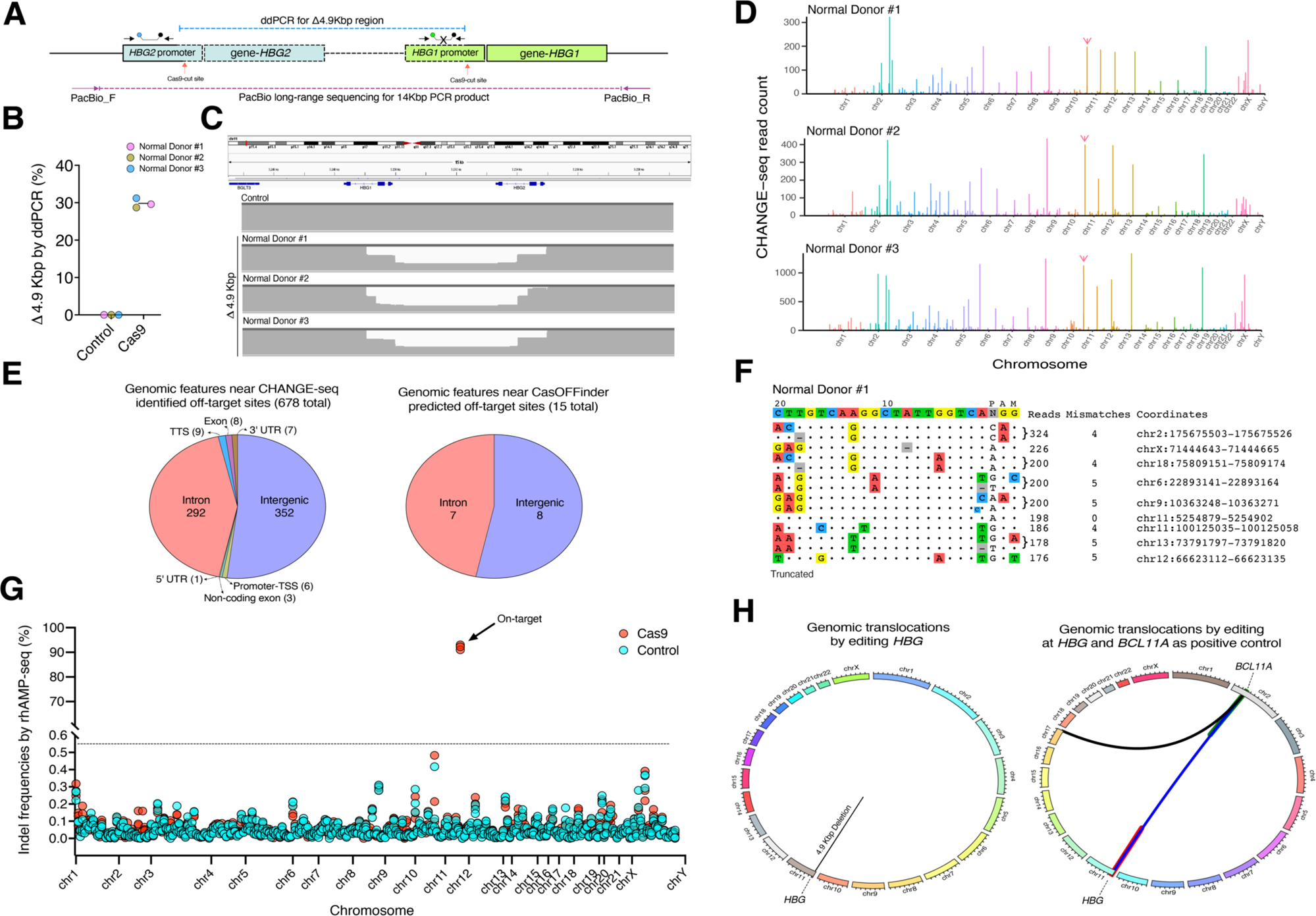
Characterization of potential on- and off-target genotoxicities associated with γ-globin promoter editing. (**A**) Schematic representation of designated ddPCR probes and primers at *HBG1* and *HBG2* gene promoters. Simultaneous Cas9-induced DNA DSBs (red arrows) results in loss of 4.9kbp deletion represented in blue dotted lines. PacBio long-range PCR primers for 14kbp region shown as purple dotted lines. (**B**) Measurement of 4.9kbp deletion by ddPCR in edited human CD34^+^ bulk HSPCs (Pre-ER#1-3, n=3, plerixafor mobilized normal donors)) at day 5. (**C**) Coverage of large deletions evaluated by long-range PCR-based PacBio sequencing in edited human HSPCs at day 5. (**D**) Manhattan plots of CHANGE-seq (n=3) detected on- and off-target sites. Intended on-target site highlighted with arrow (pink) and possible off-target sites shown as bar heights. X-axis represents chromosome location and y-axis represents number of CHANGE-seq read counts. (**E**) Alignment of intended target site (top line) with CHANGE-seq detected on- and off-target sites (top to bottom by read count) for Cas9:sgRNA (RNP) complex. Mismatches column indicates the number of mismatches relative to intended target site, where 0 indicated the on-target identified by CHANGE-seq. The coordinates column indicates the genomic coordinate, for the on- and off-target sites identified by CHANGE-seq. *Note:* output is truncated to the top sites. (**F**) Genome-wide off-target activity evaluated by multiplex targeted sequencing (rhAmp-seq, IDT) based on on- and off-target sites detected by CHANGE-seq. X-axis represents chromosomal location and y-axis indicates indels for control and Cas9 (Pre-ER#1-3, n=3) (multiple paired t-tests compared indel frequencies between edited and unedited cells using two-tailed paired t-test, controlling for false discovery rate using the procedure of Benjamini, Krieger, and Yekutieli). (**G**) Circos plots representing genomic re-arrangements for −115 γ-globin promoter and −115 γ-globin promoter + *BCL11A* enhancer as a control by UDiTaS method.

To measure the frequency of 4.9 kb deletion more precisely, we used digital droplet PCR (ddPCR) assay. Gene-specific primers and probes were designed that bind to the downstream of *HBG2* promoter and intergenic region between *HBG2* gene - *HBG1* promoter. Deletion of 4.9 kb region is estimated by loss of signal that occurs when the probe cannot bind to the intergenic deleted region compared to a control probe that binds to the *HBG2* promoter. Due to genetic variation in the binding sequences of primers and probes, it is important to check assay performance with unedited controls. We observed frequency of 29.8% (1.27) of 4.9 kb deletion in clinically scale bulk edited HSPCs (**Figure 5B**). Similar frequencies of this deletion noticed ranging from 20.8% to 37.2% in bulk edited HSPCs and 12.8% to 29.7% in the bulk BM derived from three normal and SCD donors after 17 weeks xenotransplantation (**Figure S10A**). Also, we did not detect inversions in bulk edited HSPCs and BM cells at 17 weeks post-transplantation, suggesting that these occur infrequently or that they are selected against in repopulating HSCs.

Previously, we showed by analyzing clonal burst forming unit-erythroid (BFU-E) colonies from CD34^+^ HSPCs harboring the 4.9 kb deletion, that the resultant *HBG2*-*HBG1* fusion gene retained the capacity to drive HbF expression.^33^ To further study the impact of this 4.9 kb deletion on HbF expression, we electroporated HUDEP-2 cells harboring a single intact copy of the extended β-like globin locus including the two tandem γ-globin genes^52,53^ with RNP targeting the −115 γ-globin promoter site (**Figure S11A**). The editing efficiency in bulk edited cells was 84% (1.34) and HbF was 29.4% (0.46) compared to 2.9% (0.46) in unedited control cells (**Figure S11B and S11C**). The frequency of the 4.9 kb deletion was 66% (1.38) (**Figure S11D**), higher than what typically occurs in CD34^+^ HSPCs (refer to **Figure S10A**). Analysis of edited HUDEP-2 clones revealed that with intact *HBG1* and *HBG2* genes, biallelic -13nt or -4nt deletions induced γ-globin expression 76.7% (6.52) to 78.9% (10.6) respectively. Importantly, clones with the 4.9 kb deletion and -13nt or -4nt deletions in the γ-globin fusion gene exhibited high induction of γ-globin (58.6% (6.9) to 62.7% (12.6) respectively (**Figure S11E)**, albeit to lower levels than clones with no 4.9 kb deletion and the same biallelic indels (**Figure S11F)**. Overall, these observations indicate that that erythroid cells harboring the 4.9 kb *HBG2*-*HBG1* deletion retain the capacity for strong γ-globin induction.

To characterize the spectrum of large deletions involving the extended β-globin locus that could be induced by editing the γ-globin genes, we performed long-range PCR and long-read PacBio sequencing (**Figure 5C**). We observed the major 4.9kb deletion products and no other unexpected large deletions (**Figures S12A and S12B**). One limitation of this analysis is that long-range PCR preferentially amplifies smaller DNA segments, which could overestimate the true frequency of the 4.9 kb deletion.

To define the genome-wide activity of Cas9-sgRNA directed at the γ-globin promoter genes, we used experimental and computational approaches to nominate candidate off-target sites. We performed CHANGE-seq, a highly sensitive biochemical assay to define the genome-wide off-target activity of Cas9 *in vitro*.^54^ By selective sequencing of Cas9-cleaved genomic DNA, CHANGE-seq identified 678 off-target sites in two or more replicates among from plerixafor-mobilized normal donors of African-American ancestry (pre-GMP engineering runs, n=3) (**Figures 5D and 5E**). Most of the potential off-targets classified by CHANGE-seq occurred in intergenic and intronic regions of the human genome, as expected based on their relative genomic proportions. We selected the 277 CHANGE-seq identified sites reproducibly identified in 2 or more donors and 11 additional non-overlapping CasOFFinder predicted sites for multiplexed targeted amplicon sequencing (rhAmp-seq, IDT). At day 5 post-electroporation of edited CD34+ cells mobilized from normal donors, we observed high on-target editing 92.1% (1.08) (adjusted P value: <0.000001) indels with no detectable off-target activity (FDR<0.01) from 279 amplifiable sites detected by CHANGE-seq and CasOFFinder methods (**Figure 5F**).

To assess gross genomic structural re-arrangements of edited and control cells from normal CD34^+^ HSPCs were evaluated using standard G-band karyotyping at day 5 post editing. No karyotypic abnormalities were detected in 100 metaphase spreads analyzed per edited sample (**Figure S13A**).

We applied the UDiTaS approach (Uni-Directional Targeted Sequencing)^55^ to determine the genome-wide spectrum and persistence of genomic rearrangements involving the on-target editing sites in normal donor CD34^+^ HSPCs. At day 5 and day 14 post-editing, no reproducible translocations or structural genomic re-arrangements were detected (**Figures 5G and S14A**).

In summary, through rigorous pre-clinical characterization we have found that Cas9-sgRNA complexes specifically target the −115 γ-globin gene promoters with no detectable off-target mutations and genomic re-arrangements, supporting the safety and specificity of our approach.

## Discussion

Here we described key pre-clinical studies with the goal of advancing our gene editing strategy to a future early-phase clinical trial. Typically, interactions with regulatory agencies are confidential and thus there remains no clear developmental path for IND submissions. With FDA clearance on IND based on the pre-clinical studies described here, our study may serve as a roadmap representing one path to performing IND-enabling studies responsive to recently released FDA guidance on development of human gene therapy products for genome editing.

Key criteria for advancing an ex vivo genome editing HSC cellular drug product include sufficient potency and safety predicted by pre-clinical efficacy and genotoxicity assays. Specifically, for our genome editing approach to SCD therapy, these criteria include: high HbF induction, significant reduction of sickling, broad HbF induction and no detectable genotoxicities. First, we compared different γ-globin promoter targets and identified the −115 γ-globin promoter site as our lead target site associated with the most potent induction of HbF. Second, we tested this editing strategy in mobilized, peripheral blood CD34^+^ HSPCs from individuals with SCD and confirmed high editing rates in repopulating HSCs with high induction of HbF and reduction of hypoxic sickling in erythroid progeny. Third, we performed complementary single-cell analyses that suggest broad induction of HbF containing cells. Fourth, we developed a GMP process to edit HSPCs at clinical scale using well-characterized Cas9 protein and gRNA. Analysis of the edited products showed efficient editing and robust HbF induction that is predicted to be therapeutically effective. Finally, we performed rigorous analysis of genome editing outcomes and potential unintended genotoxicities.

In pre-clinical studies, our approach appears at least comparable in potency and safety to targeting the erythroid-specific enhancer of *BCL11A*.^13^ We observed a small but significant increase in HbF induction targeting the −115 γ-globin promoter compared to the *BCL11A* enhancer target. Zeng et al. observed similar increases in their recent study. Notably, our approach also appears to be comparable in specificity with the *BCL11A-*enhancer targeting approach, with no detectable off-target activity.

Some potential advantages of our approach are its intrinsic biological specificity, straightforward path to characterize genotoxicity, and clinical scale-up. Targeting the repressor-binding site in the γ-globin promoter rather than globally depleting the expression of the repressor itself in erythroid cells may avoid any potential side effects altering non-globin *BCL11A* target gene expression.^35^ With Cas9 nuclease, we are able to utilize CHANGE-seq, a sensitive and unbiased biochemical method to characterize the genome-wide off-target activity. Towards manufacturing of patient cellular drug products, we have established GMP production of Cas9 protein and a clinical-scale process to edit cells with Cas9 RNP.

Some limitations of our Cas9 nuclease-based approach are the heterogeneity of edits that productively induce HbF^53^ and the simultaneous induction of two DNA DSBs. However, our single-cell RNA-sequencing analysis suggests that more than 80% of edited erythroblasts may induce greater than 30% of γ-globin transcripts as a proportion of all β-globin like transcripts. Our mouse studies likely underestimate the levels of HbF induction that may be achieved because beneficial anti-sickling effects of fetal hemoglobin on the half-life of sickle genotype RBCs cannot be observed in our xenotransplant experiments, because human RBCs in circulation are rapidly destroyed by mouse macrophages. A study from Frangoul et al. showed higher percentage of steady-state HbF during first-in-human trials compared to initial *in vitro* studies, which may reflect the selective advantage for HbF expressing cells. Another potential limitation is that simultaneous DSBs at the *HBG1* and *HBG2* cause deletion of 4.9kb *HBG2-HBG1* intergenic region which we detected both in the pre-transplant infusion product and engrafted BM cells. However, based on edited HUDEP2 cell clonal analysis, it was clear that cells with large deletions of *HBG2* can still be induced to express HbF through the remaining copy of *HBG1*.^56^

In sum, we describe pre-clinical characterization of a −115 γ-globin promoter editing strategy to induce HbF for treating sickle cell disease, demonstrating induction of fetal hemoglobin to therapeutically relevant levels without detectable genotoxicities. We believe this represents a promising autologous cell therapy for SCD and anticipate that our rigorous approach characterizing potential safety and efficacy could serve as a blueprint for future *ex vivo* HSC cell therapies.

## Materials and methods

### Study Design

The main purpose of this study was to demonstrate CRISPR-Cas9 genome editing of CD34^+^ HSPCs at −115 γ-globin gene promoter can induce therapeutically predictive levels of HbF for SCD therapy, to perform rigorous characterization of potential genotoxicities, and to develop a clinical-scale process towards a clinical trial. For in vitro experiments mentioned in the text, we included at least three (biological replicates) from normal donors and individuals with SCD. For *in vivo* studies, edited CD34^+^ HSPCs derived from normal and SCD donors were injected into immunodeficient NBSGW mice. Animal research technicians handling mice, injecting edited CD34^+^ HSPCs, and harvesting BM were blinded to the names of the treatment and unaware of the expected results. Experiments performed in this study with number of replicates, statistical tests, analysis, and P values are reported in figure legends and statistically significant values are mentioned in results.

### Plerixafor mobilization and CD34^+^ cells isolation

Plerixafor mobilized CD34^+^ cells were collected under approved protocol “*Peripheral Blood Stem Cell Collection for Sickle Cell Disease Patients*” (ClinicalTrials.gov identifier NCT03226691) by the human subject research Institutional Review Boards at the National Institutes of Health and St. Jude Children’s Research Hospital. All individuals gave written informed consent for the sample donation and consent documents are maintained in the donor’s medical records. The consent document was approved by the Institutional Review Board prior to study initiation and is reviewed and updated annually.

Normal plerixafor mobilized CD34^+^ HSPCs were purchased from Stem Express Stem Cell Collection Center from California. Granulocyte-Colony-Stimulating Factor (GCSF) mobilized normal HSPCs were obtained from unknown normal adults commercially purchased from Key Biologics, LLC. CD34^+^ HSPCs were isolated by AutoMACS instrument (Miltenyi Biotec) and further sorted by St. Jude Experimental Cellular Therapeutics Lab.

The process of mobilizing the CD34^+^ HSPCs from their BM niche into peripheral circulation^57^ and the details of the apheresis procedure^58^ have been described in detail previously. Donors underwent RBC exchange with goals of HbS <30% and Hb >10 g/dL prior to apheresis, received 240 µg/kg of plerixafor for HSPC mobilization, followed by HSPC collection 2-4 hours later with processing up to 4 total blood volumes. Leukapheresis was performed using a Cobe Spectra or Optia instrument via central venous apheresis catheter. Acid Citrate Dextrose formula-A (ACD-A) was used as the anticoagulant, with prophylactic IV calcium infusions used in all procedures to prevent citrate toxicity. Maximal extracorporeal blood volume during the procedure ranged from 300-400mL. We targeted deep buffy coat collection to improve SCD HSPC collection. Harvested PB apheresis products were cryopreserved using a controlled rate freezer.

### Research-grade protein purification

SpCas9-3xNLS and SpCas9-3xNLS-R691A (HF) plasmids were transformed into BL21 (DE3) competent cells (MilliporeSigma, 702353) and grown in Terrific Broth (TB) media at 37°C until reaching an OD600 2.4-2.8. Bacterial cells were induced with 0.5mM isopropyl β-d-1-thiogalactopyranoside (IPTG) for 20 hours at 20°C. Cells were pelleted and lysed in 25 mM Tris, pH 7.6, 500 mM NaCl, 5% glycerol and passed through homogenizer twice and then centrifuged at 20,000 rpm for 1 hour at 4°C. Protein was purified by Nickel-NTA resin and treated with TEV protease (1mg lab made TEV per 40mg of protein) and benzonase-100units/ml (Novagen, 70664-3) overnight at 4°C. Subsequently, purified by size exclusion column (Amersham Biosciences HiLoad 26/60 Superdex 200 17-1071-01) and ion exchange with a 5 ml SP HP column (GE 17-1151-01) according to the manufacturer’s instructions. Proteins were dialyzed in 20mM Hepes buffer pH 7.5 containing 400mM KCl, 10% glycerol, and 1mM TCEP buffer, and contaminants were removed by Toxin Sensor Chromogenic LAL Endotoxin Assay Kit (GenScript, L00350). Purified proteins were concentrated and filtered using amicon ultra-filter units (MilliporeSigma, UFC903008) and ultra-free MC centrifugal filter (MilliporeSigma, UFC30GV0S). Protein fractions were further assessed on TGX stain free 4-20% SDS-PAGE (Biorad, 5678093) and quantified by rapid gold BCA assay (Thermo Fisher, A53226).

### GMP Tag-free SpCas9-3xNLS production

A vial of *E. coli* T7 express cells containing the pGTPc502 T7 Cas9 Optimized T7-express plasmid was thawed and seeded in a 2.5L flask containing 500mL of growth media and 50μg/mL kanamycin at an OD_600_ of 0.002. Culture was incubated overnight at 30°C and 325rpm. A bioreactor (5L) was seeded at an OD_600_ of 0.4 in 2.5L of growth media (50μg/mL kanamycin) using overnight culture. Fermenter controls were 900 rpm stir speed, 30% DO, and pH 6.8 (adjusted pH with 20% NH_3_OH and 1.2 M H_3_PO_4_). FoamAway (ThermoFisher) was added as needed and air sparging was continuous at 1L/min. Reactor feed with glucose and yeast extract solution began after 5 hours of culture and increased exponentially up to 21 hours after culture start after which it remained constant. When the culture OD_600_ reached 60-80 at 20-24 hours, the temperature was decreased to 20°C and induced with IPTG (Sigma) at 0.04mM/ODU. At 21-23 hours after induction, culture was harvested by centrifugation (4000xg, 15 minutes) and the pellet was frozen at −80°C.

Three hundred grams of thawed pellet was resuspended at 10% solids in 50mM Tris (JT Baker), 350mM NaCl, 10% glycerol, pH 7.4, with 1mM TCEP (GoldBio) and 500U/g Benzonase (Millipore). Resuspended solution was lysed in a microfluidizer at 12,000 psig for two passes and centrifuged at 15,800xg for 45 minutes. Supernatant was filtered using 1040 cm^2^ 3M Harvest RC filter and passed through Sepharose SP column (BV, 461 mL) at 45cm/hr. SP Column was washed with 5 column volumes (CV) of 50mM Tris, 350mM NaCl, 10% glycerol, pH 7.4 with 1mM TCEP at 75 cm/hr followed by 8 CV elution gradient with 50mM Tris, 10% glycerol, pH 7.4, 1 mM TCEP from 350 to 750mM NaCl. Allantoin was added to Sepharose SP eluate at 100 mg/mL and stirred at 300rpm for 15 minutes at room temperature. Solution was centrifuged at 3700xg for 15 minutes and supernatant was filtered through a Sartopore 2 150 filter at 70mL/min. Next, solution was applied to a ceramic hydroxyapatite Type 1 (CHT) column (BV 132 mL) at 125 cm/hr. CHT column was washed for 5 CV with 50mM Tris, pH 8.5, 300mM NaCl, 10% glycerol, 5mM NaH_2_PO_4_, and 1mM TCEP. Elution occurred over a 20 CV linear gradient with 50mM Tris, pH 8.5, 300mM NaCl, 10% glycerol, 5mM NaH_2_PO_4_, 1mM TCEP from 300mM to 2M NaCl. The eluted fractions were combined and concentrated to 15-20 mL using a pellicon mini 0.1 cm^2^ biomax 100 kDa cutoff membrane at 80mL/min and diafiltrated with 20 CV of 20mM HEPES, 300 mM KCl, 10% glycerol, pH 7.4. Final concentration measured by UV spectroscopy and protein was diluted to 12mg/mL with 20 mM HEPES, 300mM KCl, 10% glycerol, pH 7.4 before final filtration using a 0.22μM millex syringe filter.

### Cell culture

CD34^+^ HSPCs were thawed and pre-stimulated for 48 hours in X-VIVO 15 (Lonza, 04-418Q), and for pre-clinical and clinical-scale up experiments we used X-VIVO 10 (Lonza, 04-743Q) supplemented with 100 ng/mL of SCF, Flt3-L, TPO, and 1% penicillin/streptomycin. SCD derived CD34^+^ HSPCs were pre-stimulated for 24 hours prior to EP. Cells were washed with 1xPBS, counted, and resuspended in P3 solution. Electroporated cells were recovered in X-VIVO media with SCF, Flt3-L, and TPO at 37°C, 5% CO2. After 24 hours, cells were cryopreserved in media containing 0.5ml of plasma-Lyste A + 4% HSA and 0.5ml of 2x Freezing medium includes final concentrations of 12% pentastarch, (20% USP, Preservation Solutions Inc, PST001), 10% DMSO, 7.5% HSA. For erythroid differentiation, post-electroporated cells were transferred into phase 1 media (1-7 days) with Iscove’s modified Dulbecco’s medium (IMDM) containing cytokines and penicillin/streptomycin, followed by culturing in phase 2 (8-14 days), and phase 3 (15-21 days). Phase 1 consists of IMDM (Gibco, 12440053) with 2% human blood type AB plasma (SeraCare, 1810-0001), 3% human AB serum (Atlanta Biologicals, S40110), 1% penicillin/streptomycin (Gibco, #15140122), 3 units/mL heparin (Sagent Pharmaceuticals, NDC 25021-401-02), 3 units/mL EPO (Amgen, EPOGEN NDC 55513-144-01), 200 μg/mL holo-transferrin (Millipore Sigma, T0665, 10 ng/mL recombinant human SCF (R&D systems, 255-SC/CF), and 1 ng/mL recombinant human interleukin IL-3 (R&D systems, 203-IL/CF). Phase 2 includes phase 1 media without IL-3 and phase 3 includes phase 1 media without IL-3 and SCF with holo-transferrin increased to 1 mg/mL final concentration. Cells were maintained at 1.0-2.0 x 10^5^ cell/mL in phase 1 and phase 2, whereas phase 3 seeding density is at 1.0-2.0 x 10^6^ cell/mL. At day 21 in phase 3 media, fully differentiated cells were collected for flow cytometry and HPLC. Non-electroporated cells are included as controls.

### Genome editing

Cas9-3xNLS and Cas9-3xNLS-R691A (HF) proteins were purified by St. Jude production core facility and tag-free Cas9-3xNLS purified by St. Jude GMP facility. Synthetic sgRNAs (chemically modified with 2’-O-methyl analogs and 3’ phosphorothioate internucleotide linkages at the first three 5’ and 3’ terminal RNA residues) were ordered from Synthego (**Table S1**). Lyophilized gRNAs were reconstituted in nuclease-free water and aliquots were stored at −80°c. Electroporation performed using Lonza 4D Nucleofector with P3 Primary Cell Nucleofector Kit (V4SP-3096) for 20 µL and (V4XP-3024) for 100µL reaction. Cas9-RNPs complexes were made by combining 50 pmoles of SpCas9-3x-NLS and 150 pmoles sgRNA kept for 15 mins at room temperature to electroporate 2 x 10^5^ cells in 20 µL reaction and 250 pmoles of SpCas9-3x-NLS, 750 pmoles of sgRNA were used to nucleofect 4 x 10^6^ cells in 100 µL P3 reaction. Optimal programs for Cas9-3xNLS and Cas9-3xNLS (HF) using Lonza 4D nucleofector is DS-130 and CM-137. Electroporated cells were recovered in warmed media for 5 mins and cultured in X-VIVO media with cytokines at 37°C, 5% CO2. Cells were harvested for NGS at day 4 or 5.

### Optimization and clinical scale up electroporation

For optimization and scale up, cells were electroporated using the MaxCyte GTx. Apheresis of GCSF or plerixafor mobilized donors were shipped overnight and purified for CD34^+^ cells. Where cells were processed according to manufacturer’s instructions with the Sepax C-pro and then the CliniMACS Plus with CD34^+^ selection beads. The cells were then either cryopreserved or pre-stimulated overnight in X-VIVO 10 containing 100 ng/mL of SCF, Flt3-L, and TPO. For optimization studies, we used the JMP (SAS) software to conduct DOE studies looking at key parameters for the editing of the HSCs, which include Cas9 concentration, Cas9 to sgRNA ratio, cell concentration, with key outcomes including editing efficiency, cell viability, and cell recovery. After pre-stimulation between 0 to 48 hours, the cells were then washed with MaxCyte buffer with 0.1% HSA. The cells were then resuspended with Maxcyte buffer between 20-200 million cells/mL. The RNP was formulated between 1 to 9.6uM of Cas9 protein at 1 to 3 molar ratio of sgRNA. The cells and the RNP were combined and mixed and transferred to an OC-100 or CL1.1 cassette and electroporated with programs HSC3, HSC4, or HSC5. After EP the cells were transferred to fresh X-VIVO 10 with cytokines at 2E^6^ cells/mL and cultured overnight. The cells were then cryopreserved at 5×10^6^ cells/mL in a freezing medium containing Plasma-Lyte A with 5% dimethyl sulfoxide (DMSO), 6% pentastarch, and 5.75% human serum albumin (HSA) with a controlled rated freezer. The final large-scale process was finalized with pre-stimulation of HSCs before EP for 24 hours and cryopreservation after 24 hours EP. EP conditions was locked in at 4.8mM of Cas9 with 1:2 Cas9 to sgRNA ratio with EP of cells between 10-100 million cells per mL in a CL1.1 cassette with program HSC3.

### High-throughput sequencing

Genomic DNA was isolated using DNAdvance Kit (Agencourt, A48705). Targeted sequencing libraries for NGS were prepared using a 2-step PCR protocol. Step one creates targeted sequencing amplicons using gene specific primers containing a partial illumina adaptor overhang (**Table S2**). PCR amplifications were performed using phusion hot start flex 2X master mix (NEB, M0536S) and 20-100ng genomic DNA with the following conditions for HBG-sgRNA target (denaturation - 98°C for 3:00; touchdown (20x cycles) - 98°C 68°C 72°C for 0:10 0:10 (0.5°/cycle) 0:07; annealing (10x cycles) - 98°C 58°C 72°C for 0:10 0:10 0:07; final extension 72°C for 3:00; hold 10°C. PCR conditions for *BCL11A* enhancer or ZBTB7A are denaturation - 98°C for 3:00; annealing (35x cycles) - 98°C 60°C 72°C for 0:15 0:15 0:30; final extension 72°C for 3:00; hold 10°C. PCR step one product was purified using AMPureXP beads. PCR step 2 was conducted using 10-15ng PCR1 product to complete Illumina sequencing adaptors (P5-dual-index. F and P7-dual-index. R) using Kapa HF Hot Start Ready Mix (Roche, KK2602) with following PCR conditions denaturation - 98°C for 0:45; annealing (10x cycles) - 98°C 65°C 72°C for 0:15 0:30 0:30; final extension 72°C for 1:00; hold 10°C. PCR2 products were purified with AmpureXP beads, pooled each sample with 1000nM and further diluted to 1.5nM in 1xTE. Library was sequenced using Miseq (Illumina) and Miseq micro v2 kit and sequenced following manufacturers protocol spiked with 10% Phix control (FC-110-3001). Library was sequenced using paired end sequencing with 151-8-8-151 cycles.

### Transplantation of cells into NBSGW mice

Nonirradiated NOD, B6.SCID Il2rγ−/−KitW41/W41 (NBSGW) (stock no. 026622) were purchased from the Jackson Laboratory. 5 x 10^5^ cells (control or edited) administered by tail-vein injection to 8-12 weeks of female mice. 17 weeks post transplantation, BM was harvested and evaluated for donor chimerism and indels. CD235a^+^ erythroblasts were isolated from bulk BM with magnetic beads, using the human specific CD235a^+^ glycophorin A MicroBeads (Miltenyi Biotec Inc, 130-050-501). Purified CD235a^+^ were evaluated for erythroid maturation and enucleation (**Table S3)**. Harvested BM cells were sorted using FACSAria III cell sorter (BD Biosciences) to isolate various cell lineages (hCD45, hCD3, hCD19, hCD34, hCD235a) using human lineage specific antibodies (**Table S4**).

### In vitro sickling assay

CD34^+^ HSPCs were differentiated as mentioned above using three-phase protocol. Erythroid cells in phase 3 media were incubated with Hoechst 33342 (Millipore Sigma) at 1:1000 dilution for 20 mins at 37°C and sorted for hoechst-negative population using a SH800 (Sony Biotechnologies). 96-well plate pre-coated with poly-L-lysine (Sigma) for 30mins and washed with 1xPBS. 5 x 10^4^ sorted cells were seeded into 96-well plate with 0.1 ml of phase 3 medium under hypoxic conditions (2% oxygen) for 8 hours. The IncuCyte S3 Live-Cell Analysis System (Sartorius) with a 20x objective was used to monitor cell sickling, with images captured after 8 hours. The percentage of sickling was measured by manual counting of sickled cells versus normal cells based on morphology. For each sickling assay, more than 300 cells per condition were counted by researchers blinded to that condition.

### Single Cell Western

MACS purified erythroid (CD235a^+^) cells from BM were pooled and 8 x 10^4^ cells were loaded on small scWest chips (Protein Simple, Cat # K500). scWest chips were processed with lysis time for 0 sec, electrophoresis for 30 sec, voltage at 240 V, and UV exposure for 4 mins according to the manufacturer’s protocol. Probed with primary antibodies rabbit anti α-globin from ResGen (Invitrogen Corporation) and mouse anti γ-globin (Santa Cruz, Sc-21756, 51-7) at dilutions 1:1500 and 1:10 followed by secondary antibodies Donkey anti-rabbit IgG NorthernLights557 (R&D Systems, NL004) and Donkey anti-mouse-Alexa Fluor647 (Fischer, A31571) at 1:40 dilution. scWest chips were scanned by InnoScan710 micro array scanner and 16-bit TIFF images were analyzed by scout software.

### Single cell RNA sequencing (ScRNA-seq)

Pooled erythroid (CD235a^+^) cells were used for ScRNA-seq studies. Determined cell count and appropriate volume of cells was loaded on the Chromium Next GEM chip G to run on Chromium controller according to the Chromium Next GEM Single Cell 3’ reagent kits v3.1 user guide (Cat#1000127, 1000128) for a targeted recovery of 9000 cells. Post GEM-RT clean up, cDNA amplification, and library construction was performed according to the 10X Genomics Next GEM single cell protocol and Agilent 2100 Bioanalyzer was used for cDNA QC, quantification, and post library construction QC. Sequenced by using Novaseq6000 with library concentration 1.3nM (+1% Phix) loaded on S1 flow cell and read length for R1-i7-i5-R2 was 28-8-0-91 cycles. ScRNA-seq data was analyzed using cellranger count (v7.0.0) with default parameters. The transcriptome database was based on cellranger GRCh38-2020-A_build. Since the cellranger algorithm only counted reads that were uniquely assigned to a single gene, to get accurate quantification of *HBG* genes, two novel transcripts in the database (ENSG00000284931 and ENSG00000239920) that overlapped with *HBG1* and *HBG2* genes were removed. Seurat v4.3.0 was used for the downstream analysis. The single-cell data were filtered by: (1) number of expressed genes (i.e., nFeature_RNA) between 200-6000, (2) percentage of mitochondrial reads (i.e., percent.mt) less than 10% and (3) the total read (i.e., nCount_RNA) less than 50000. The proportion of HBG gene expression was computed as the total read count of *HBG1* and *HBG2* divided by the total read count of *HBG1*, *HBG2*, and *HBB* genes. The “FindMarkers” function was used for differential gene expression analysis comparing edited and unedited cells. Single-cell data integration was based on the “anchor” method. The UMAP (Uniform Manifold Approximation and Projection) plot was generated using the first 50 PCs. Cell clustering was performed on the UMAP space with resolution of 0.1. Cell annotation was based on gene markers identified in ^45^.

### Single guide RNA (sgRNA) sequencing and analysis

Synthetic gRNAs (1-2ng) were sequenced by SMARTer smRNA-seq kit from Takara (Cat #635030). Original oligo-dT protocol is modified and first cDNA was synthesized directly with 1µl of tailed scaffold primer (10µM) that specifically binds to gRNA scaffold sequence 5’ GTGACTGGAGTTCAGACGTGTGCTCTTCCGATCT <u>TGCCACTTTTTCAAGTTG</u> 3’. Incubate the reaction at 72°C for 3 mins and cool to 4°C for 2 mins to anneal the primer at the 3’ end of the scaffold gRNA. Reverse transcription master mix added to the reaction according to the manufacturer’s instructions. Next, full-length illumina adapters were added by PCR primers with the following conditions, denaturation - 98°C for 1:00; annealing (17x cycles) - 98°C 60°C 68°C for 0:10 0:05, 0:10; hold 10°C. PCR products were purified by nucleospin clean up kit and quantified by Qubit HS. Pooled index library 2.2nM (+10% Phix) sequenced on MiSeq (Illumina) with MiSeq v2 micro kit using paired end sequencing with 101-8-8-101 cycles. Paired-end reads were merged using FLASH with an expected fragment length of 100bp. Scaffold specific primer (3’-end) was trimmed using cutadapt (v4.1) and read with length than 20bp were removed. The percentage of target gRNA and full-length product (target gRNA + scaffold sequence) were based on an exact string match. Merged reads were aligned to the expected full-length product using bowtie2 to identify indels in the gRNA sequencing data. Source code is available at: https://github.com/YichaoOU/gRNA_sequencing.

### Digital droplet PCR (ddPCR) for 4.9-kb deletion

Specific probes were designed to target *HBG1* and *HBG2* regions and used with respective forward and reverse primers that amplify both *HBG1* and *HBG2,* resulting in 198bp and 106bp amplicons. Alternate primer set were designed for donor specific regions with nucleotide changes (**Table S5**). The reaction was set up with 100-300ng of genomic DNA (from bulk BM), 0.9μM probes, 0.25μM probes, and ddPCR supermix (no dUTPs) (Bio-Rad, 186304); droplets were generated using automated droplet generator (Bio-Rad auto DG) and PCR reaction was performed on a CX1000 thermo cycler with the following steps: 95°C for 10 min; 94°C for 30 sec; 56°C for 1 min (ramp rate 2°C/sec, go to step 2 for 40 cycles total); 98°C for 10 min; 4°c hold. Droplets were read by using QX200 droplet reader (Bio-Rad) to measure the fluorescence intensity and data were analyzed using QuantaSoft.

### PacBio sequencing

200ng of genomic DNA used isolated from DNAdvance Kit (Agencourt, A48705) to PCR 14.8kb region in the *HBB* locus using forward and reverse PCR primers with indexes to allow pooling of multiple donor samples (**Table S5**). PCR conditions are denaturation - 98°C for 0:30; annealing (25x cycles) - 98°C 68°C 72°C for 0:10 0:20 7mins; final extension 72°C for 10:00; hold 4°C. The barcoded PCR reactions were cleaned using 1X Pacific Bioscience AMPure PB beads before being quantified via picogreen (ThermoFisher) and sized via Agilent Femto Pulse 165Kb gDNA assay. The cleaned PCR products were then pooled, and sequence ready libraries were generated using Pacific Biosciences SMRTbell Express Template Prep Kit 2.0. Sequencing was performed on a Pacific Biosciences Sequel IIe using sequencing primer v5, Sequel II binding kit 2.2, DNA internal control v1.0, using adaptive loading and pre-extension. Sequencing results were analyzed by SMRTtools (v10.1) from PacBio was used to convert bam files to fastq files. Then the reads were aligned to the human genome (hg38) using minimap2 with options of “-ax map-hifi --MD --secondary=no”. Indels were extracted from the resulting sam file.

### HbF measurement

Analytical ion-exchange high performance liquid chromatography (IE-HPLC) quantification of hemoglobin tetramers and individual globin chains was performed using ion-exchange columns on a Prominence HPLC System (Shimadzu Corporation). Proteins eluted from the column were identified at 220 and 415 nm with a diode array detector. The relative amounts of hemoglobin’s or individual globin chains were calculated from the area under the 415-nm peak and normalized based on the DMSO control. Percentage of γ-globin = ((Gγ-chain + Aγ-chain)/β-like chains (β + Gγ + Aγ) × 100.

### CHANGE-seq off-target analysis

Genomic DNA from human CD34^+^ cells from three normal adult donors was isolated using Gentra Puregene Kit (Qiagen) according to manufacturer’s instructions. CHANGE-seq was performed as previously described ^54^. Briefly, purified genomic DNA was tagmented with a custom Tn5-transposome to an average length of 400 bp, followed by gap repair with Kapa HiFi HotStart Uracil+ DNA Polymerase (KAPA Biosystems) and Taq DNA ligase (NEB). Gap-repaired tagmented DNA was treated with USER enzyme (NEB) and T4 polynucleotide kinase (NEB). Intramolecular circularization of the DNA was performed with T4 DNA ligase (NEB) and residual linear DNA was degraded by a cocktail of exonucleases containing Plasmid-Safe ATP-dependent DNase (Lucigen), Lambda exonuclease (NEB) and Exonuclease I (NEB). *In vitro* cleavage reactions were performed with 125 ng of exonuclease-treated circularized DNA, 90 nM of SpCas9-3x-NLS protein, NEB buffer 3.1 (NEB) and 270 nM of HBG-sgRNA (Synthego), in a 50 μL volume. Cleaved products were A-tailed, ligated with a hairpin adaptor (NEB), treated with USER enzyme (NEB) and amplified by PCR with barcoded universal primers NEBNext Multiplex Oligos for Illumina (NEB), using Kapa HiFi Polymerase (KAPA Biosystems). Libraries were quantified by qPCR (KAPA Biosystems) and sequenced with 151 bp paired end reads on an Illumina NextSeq instrument. CHANGE-seq data analyses were performed using open-source CHANGE-seq analysis software (https://github.com/tsailabSJ/changeseq).

### Targeted amplicon sequencing by rhAmpSeq

On- and off-target sites identified by CHANGE-seq and CasOFFinder were amplified from genomic DNA from edited CD34^+^ normal cells or unedited control donor cells using rhAmpSeq system (IDT). Sequencing libraries were generated according to the manufacturer’s instructions and sequenced with 151-bp paired-end reads on an Illumina NextSeq instrument.

### Sequencing analysis

Indel sequencing or rhAmp-seq analysis was conducted using custom Python code and open-source bioinformatic tools. First, high-throughput sequencing data files were demultiplexed to generate FASTQ format files. Paired-end high-throughput sequencing reads were processed to remove adapter sequences with trimmomatic ^59^ (version 0.36), merged into a single read with FLASH ^60^ (version 1.2.11) and mapped to human genome reference hg38 using BWA-MEM ^61^ (version 0.7.12). Reads that mapped to on-target or off-target sites were realigned to the intended amplicon region sequence using a striped Smith-Waterman algorithm as implemented in the Python library scikit-bio; indels were counted and reported with total read counts.

Targeted amplicon sequencing data was analyzed using CrisprEsso (v2.0.41). Microhomology-mediated end joining (MMEJ) was annotated based on sequence homology between the deleted sequence and its flanking upstream and/or downstream sequence. Specifically, the following two cases were labeled as MMEJ: if (1) the first 2bp of the deleted sequence was the same as the first 2bp of its downstream sequence or (2) the last 2bp of the deleted sequence was the same as the last 2bp of its upstream sequence.

### Statistics

Statistical analyses were performed using GraphPad Prism 10.0 (GraphPad Software Inc., La Jolla, CA). For comparative analysis ordinary one-way ANOVA (Dunnett’s multiple comparison tests and Tukey’s multiple comparison tests) and two-way ANOVA (Šídák’s multiple comparisons tests) were used. Unpaired t-tests were performed for comparison for fewer than two groups. Data points with three or more replicates considered for statistical analyses and two or fewer were excluded. For mean and SEM, row statistics was performed.

Figure 4A created with BioRender.com

**Supplementary Data** is available in the online version of the original paper.

## Acknowledgements

We thank the individuals with sickle cell disease who contributed to the study and following core facilities and individuals at St. Jude Children’s Research Hospital: Flow Cytometry (Richard Ashmun and Stacie Woolard), Animal Resource Center (Chandra Savage), Protein production core (Richard Heath), Hartwell Center (Geoffrey Neale), and Cytogenetic Core (Marc Valentine). Hemizygous HUDEP2 cells (*HUDEP2*^Δυγδβ^) were generated and kindly shared by Phillip Doerfler. Allophycocyanin (APC)-conjugated anti-Band3 was a gift from X. An (New York Blood Center).

## Funding

This work was supported by the Doris Duke Charitable Foundation (DDCF) grant (2017093 and 2020154 for aspects of this study that did not utilize mice to S.Q.T. and M.W.), National Institutes of Health (NIH) Office Of The Director (OD) Somatic Cell Genome Editing (SCGE) initiative grants U01AI157189 and U01AI176471 (to S.Q.T.), St. Jude Children’s Research Hospital Collaborative Research Consortium on Novel Gene Therapies for Sickle Cell Disease (SCD), and St. Jude Children’s Research Hospital and ALSAC. St. Jude Children’s Research Hospital core facilities are supported by NIH grant P30CA21765. DBT-Ramalingaswami re-entry fellowship, Govt. of India (to T.M). The article is solely obligated to the authors and does not necessarily represent the official views of the funders such as NIH or the DDCF. The funders had no role in study design, data collection and analysis, decision to publish or preparation of the manuscript.

## Author contributions

V.K., K.O.K, Y.L., T.M., C.R.L., R.W., G.L., Y.Y., R.L., K.M., Y.J., N.N., E.D., S.Z., contributed to experiments, data collection and data analyses. A.P., T.L., M.M., F.F contributed to GMP Cas9 protein production and characterization studies. A.S., N.U., J.F.T., provided CD34^+^ HSPCs from patients. V.K., S.Q.T., J.S.Y., M.J.W drafted and edited manuscript. M.J.W and S.Q.T. conceived of the study design, supervised the study, and revised the final manuscript.

## Conflicts of Interest

A.S. has received consultant fee from Spotlight Therapeutics, Medexus Inc., Vertex Pharmaceuticals, Sangamo Therapeutics and Editas Medicine. He is a medical monitor for RCI BMT CSIDE clinical trial for which receives financial compensation. He has also received research funding from CRISPR Therapeutics and honoraria from Vindico Medical Education.

A.S. is the St. Jude Children’s Research Hospital site principal investigator of clinical trials for genome editing of sickle cell disease sponsored by Vertex Pharmaceuticals/CRISPR Therapeutics (NCT03745287), Novartis Pharmaceuticals (NCT04443907) and Beam Therapeutics (NCT05456880). The industry sponsors provide funding for the clinical trial, which includes salary support paid to A.S.’ institution. A.S. has no direct financial interest in these therapies.

J.S.Y is an equity owner of Beam Therapeutics.

M.J.W is on advisory boards for Cellarity Inc., Novartis, and Forma Therapeutics.

S.Q.T is a co-inventor on licensed patents for CHANGE-seq and other genome engineering technologies. S.Q.T. is a member of the scientific advisory boards of Prime Medicine and Ensoma.

Other authors declare no competing non-financial interest.

## Supplemental Methods

### Erythroid culture and flow cytometry

CD34+ HSPCs were differentiated into erythroid cells in vitro using a three-phase protocol ^1^. Erythroid maturation during in vitro differentiation was measured by immuno-flow cytometry for the cell surface markers CD235a, CD49d and Band3 (**Table S4**). Erythroid enucleation rate and F-cell population analysis: 1.0−2.0 × 10^5^ CD34^+^ HSPC-derived erythroid cells were incubated with Hoechst 33342 for 20 min at 37°C, fixed with 0.05% glutaraldehyde (Millipore Sigma, G5882), and permeabilized with 0.1% Triton X-100 (Millipore Sigma, 93443). Subsequently, cells were stained with CD235a and anti-human HbF then analyzed by flow cytometry.

### MLPA assay

Genomic DNA extracted from bulk BM cryopreserved cells following manufacturer’s protocol from cultured cells using gentra puregene kit (Qiagen, 155867). 100-200ng of genomic DNA was used to perform denaturation, hybridization, ligation, and PCR according to the manufacturer’s instructions using SALSA-MLPA probe kit (Cat # P102-D1 HBB, MRC-Holland). PCR fragments were separated by capillary electrophoresis using ABI 3730xl (G5 set filter & GS500 LIZ) and MLPA data were analyzed using Cofalyser.Net (MRC-Holland).

### HUDEP-2 culture

Hemizygous HUDEP-2 cells (HUDEP2^Δεγδβ^) were expanded in culture medium consisting of SFEM (Stem Cell Technologies, catalog no. 09650) supplemented with 50 ng/ml-1 recombinant human SCF, 3 IU ml-1 EPO, 1 μg/ml-1 doxycycline (Sigma Aldrich, catalog no. D9891), 0.4 μg/ml-1 dexamethasone (Sigma Aldrich, catalog no. D4902), and 1% penicillin-streptomycin solution. HUDEP-2 cells were differentiated using a two-phase protocol. In phase 1 (days 1-3), cells were maintained in IMDM with 2% fetal bovine serum, 2% human blood type AB plasma, 1% penicillin/streptomycin, 3 units/ml-1 heparin, 10 μg/ml-1 insulin (Sigma, catalog no. I9278), 3 units/ml-1 EPO, 1 mg/ml-1 holo-transferrin, 50 ng/ml-1 SCF and 1 µg/ml-1 doxycycline. In phase 2 (days 5–10), cells were maintained in phase 1 medium without SCF.

Hemizygous HUDEP2 cells were electroporated with Cas9 using Lonza 4D nucleofector system with the DS-150 program. Edited hemizygous HUDEP2 cells were sorted into 96-well round-bottom plates (Corning) at one cell per well in 100 μl of expansion medium, using SH800 sorter (Sony Biotechnologies). The clones were expanded for 14 days and later differentiated using the two-phase culture protocol, and the percentage HbF was measured by IE- and RP-HPLC.

HbF quantification session: Percentage of γ-globin hemoglobin subunits was calculated from reverse-phase HPLC as follows: percentage γ-globin = ((Gγ-chain + A γ-chain)/β-like chains (β + Gγ + Aγ)) × 100.

### Measurement of *HBG2* loss by NGS

The *HBG2* loss in the HUDEP-2 clones were measured by NGS as described previously ^2^. Briefly, oligonucleotide primers (see below) were used to amplify exon 3 of both *HBG1* and *HBG2* to generate 261 bp gene-specific amplicons that differ by 1 nucleotide at cDNA position 410 (G or C at *HBG2* and *HBG1*, respectively) and analyzed by targeted NGS, with reduced G reads in the common amplicon reflecting the deletion of *HBG2* exon 3.

Underlined sequences correspond to universal 5’ tails.

ΔHBG2.F CACTCTTTCCCTACACGACGCTCTTCCGATCTGGTGGAAGCTTGGTGTGTAG ΔHBG2.R GTGACTGGAGTTCAGACGTGTGCTCTTCCGATCTAAAGCCTATCCTTGAAAGCTCT

### Capillary based western blot

EZ standard reagents were prepared 400mM DTT, 5x fluorescent master mix, and biotinylated ladder following manufacturer’s protocol. 250k cells were lysed using 65-85µl of 1x RIPA buffer (Millipore Sigma, R2078) with protease inhibitor cocktail (Sigma, P8340, 500X). Kept on ice for 15mins, centrifuge at maximum speed for 15mins at 4°C and saved supernatant (−80°C). Denature 4µl of supernatant with 1µl of 5x fluorescent master mix at 95°C for 5mins and kept on ice. Primary antibodies rabbit anti Cas9 pAb (Takara, 632607) and mouse anti beta-actin mAb (Sigma, A2228) were diluted in the same tube of milk free diluent at 1:20 and 1:30 dilution. Secondary antibodies goat anti rabbit NIR and goat anti mouse IR from kit were diluted at 1:20 in milk free diluent. Dispense prepared samples into 12-230kDa cartridge (Protein Simple, AM-PN01) and run jess according to manufacturer’s instructions.

### Karyotyping

Human CD34^+^ HSPCs derived from GCSF mobilized normal donors and edited at −115 g-globin gene promoters. At Day 5, cells were harvested after a four-hour colcemid exposure for gross karyotype analysis (G-banding). For standard karyotyping, cells were air dried on slides with trypsin and Wright’s stain using traditional cytogenetic methods. Twenty metaphase cells were characterized per treatment condition. Unedited cells were used as a control.

### Analysis of translocations and other structural rearrangements

Unidirectional targeted sequencing (UDiTaS) ^3^ protocol performed to identify translocations with the on-target breakpoint of HBG-sgRNA in the HBG1/HBG2 promoter at days 5 and 14 after editing. A custom Tn5-transposome with the full-length Illumina forward (i5) adapter, a sample barcode, and unique molecule identifier (UMI) was used for the genomic DNA tagmentation reaction. After genomic DNA tagmentation, two rounds of PCRs were performed. The first PCR was performed using the target specific HBG promoter anchor primer **(Table S7**), and the second round of PCR added the reverse (i7) Illumina adapter and an additional sample barcode. Two primers targeting HBG1 and HBG2 promoter were used in parallel for the first PCR. Conditions for PCR1 and PCR2 are denaturation - 95°C for 5:00; touchdown (15x cycles) - 95°C 70°C 72°C for 0:30 2:00 (1°C /cycle) 0:30; annealing (10x cycles) 95°C 55°C 72°C for 0:30 1:00 0:30; 72°C for 5:00; hold 4°C. Complete libraries were then size selected for 400-850bp using PippinHT, quantified by qPCR and sequenced on Illumina MiSeq with paired end 301-8-18-301 cycles. Analysis performed by (https://github.com/editasmedicine/uditas).

## Supplemental Figures

**Figure S1:**
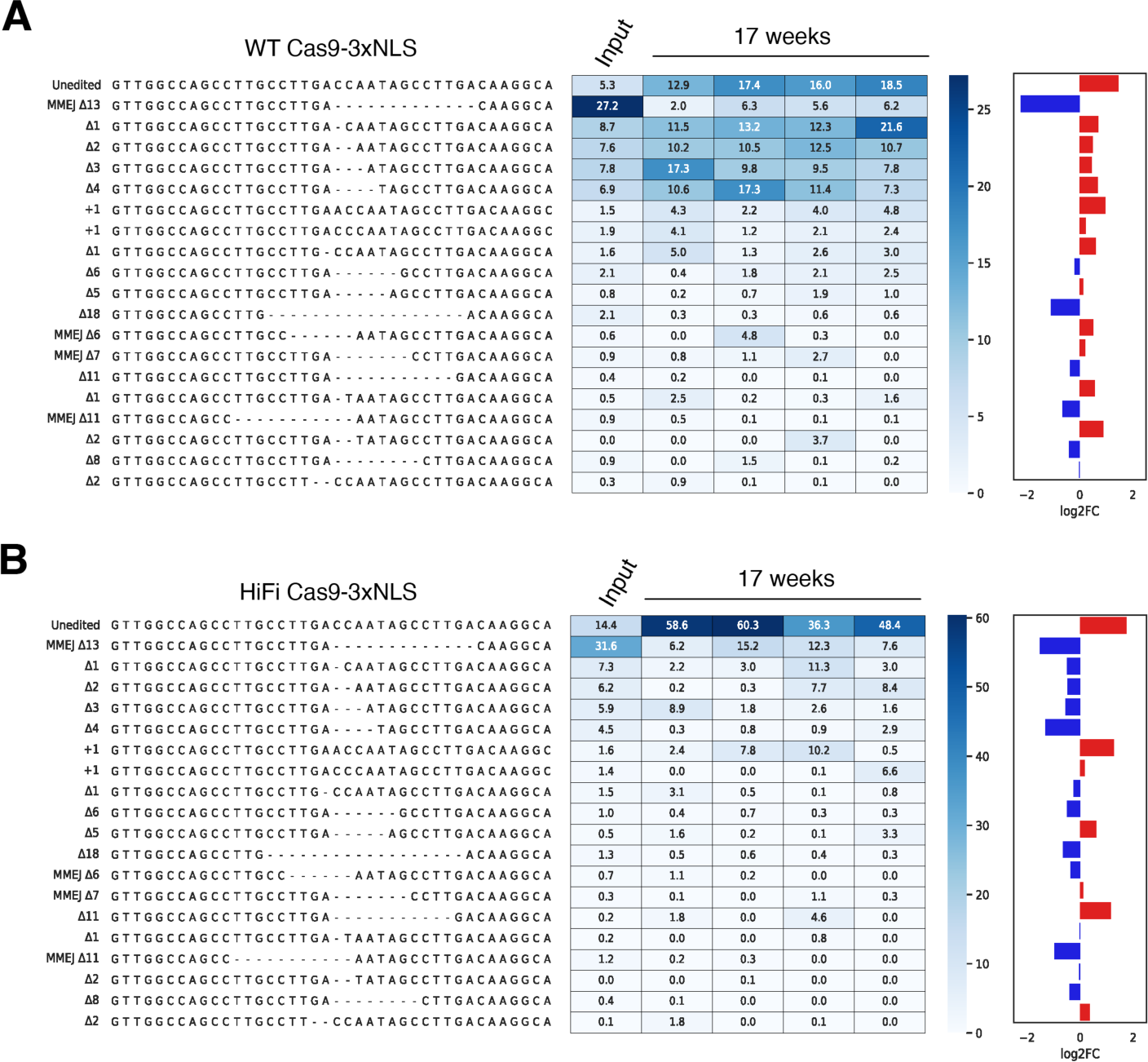
Comparison of indels spectrum with wild-type and high-fidelity Cas9-3xNLS at −115 g-globin gene promoters after 17 weeks transplantation. Representation of common editing outcomes caused by WT (A) and HiFi (B) Cas9-3xNLS due to disruption −115 g-globin gene promoters. Length of the deletions and insertions shown on left panel and mutations related to microhomology-mediated end joining pathway annotated as MMEJ. Heat map showing indel frequency (%) in CD34+HSPCs (input) prior to xenotransplantation and repopulating human HSCs after 17 weeks of engraftment in NBSGW mice (n = 4). Bar plot showing log2 fold changes of indel frequency, 17 weeks (red) vs input (blue).

**Figure S2:**
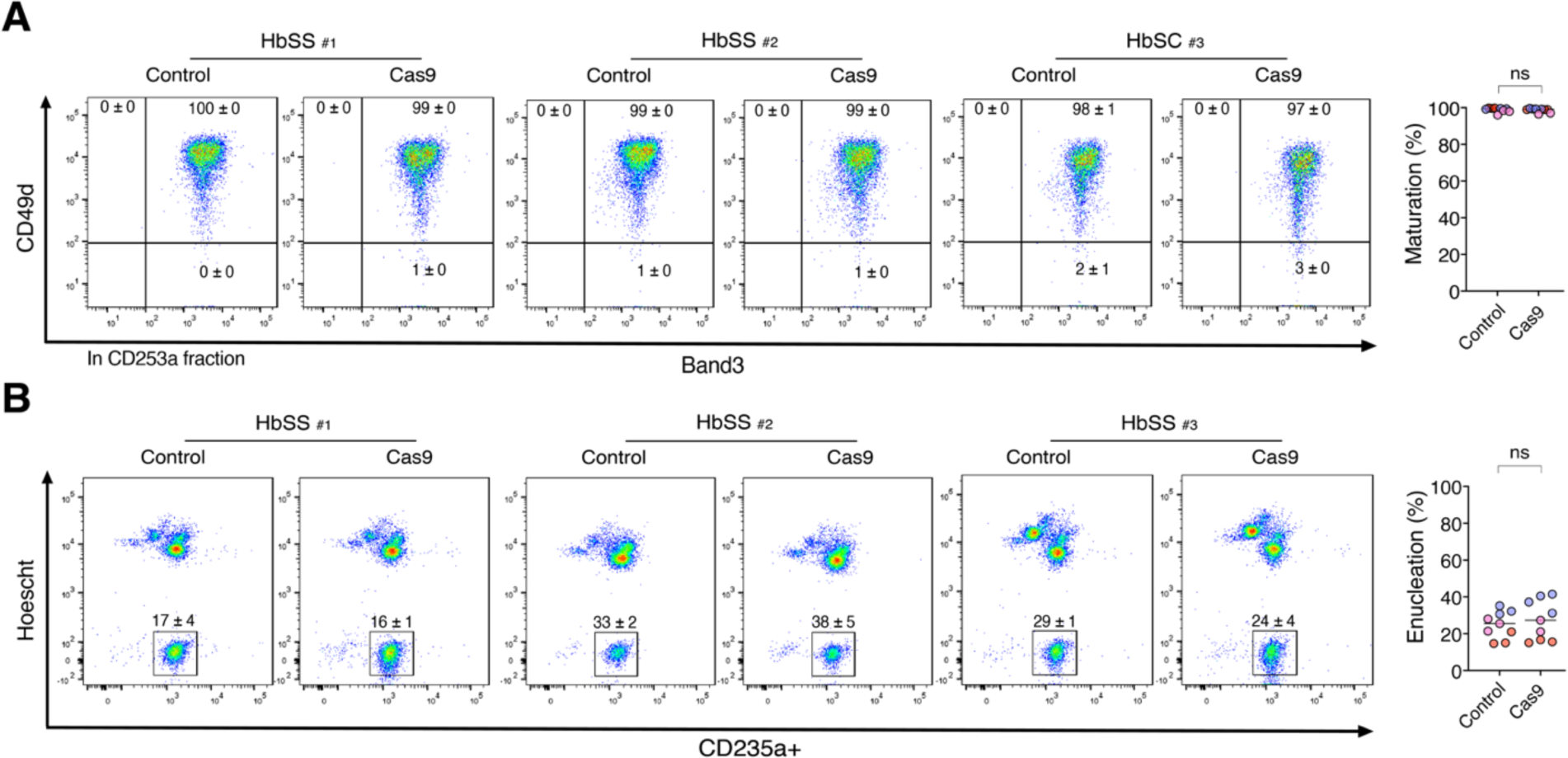
Erythroid maturation and enucleation of edited human HSPCs derived from three SCD donors in transplanted NBSGW mice. (**A**) Measurement of erythroid maturation in human CD235a^+^ fractions with antibodies against CD49d (early erythroblast marker, shown on y-axis) and Band3 (late maturation marker, shown on x-axis). On right, dot plot shows the percentages of maturation for erythroid cells for control and Cas9. (Unpaired t-test, ‘ns’ not-significant, P value 0.7627) (**B**) Enucleation of erythroid cells was examined with hoechst staining (nuclear stain, shown on y-axis) and CD235a^+^ (shown on x-axis) by flow cytometry. Statistically analysis indicates no significant changes between control and Cas9 edited cells. Percentage of enucleation for erythroid cells for control and Cas9 (Unpaired t-test, ‘ns’ not-significant, P value 0.5729).

**Figure S3:**
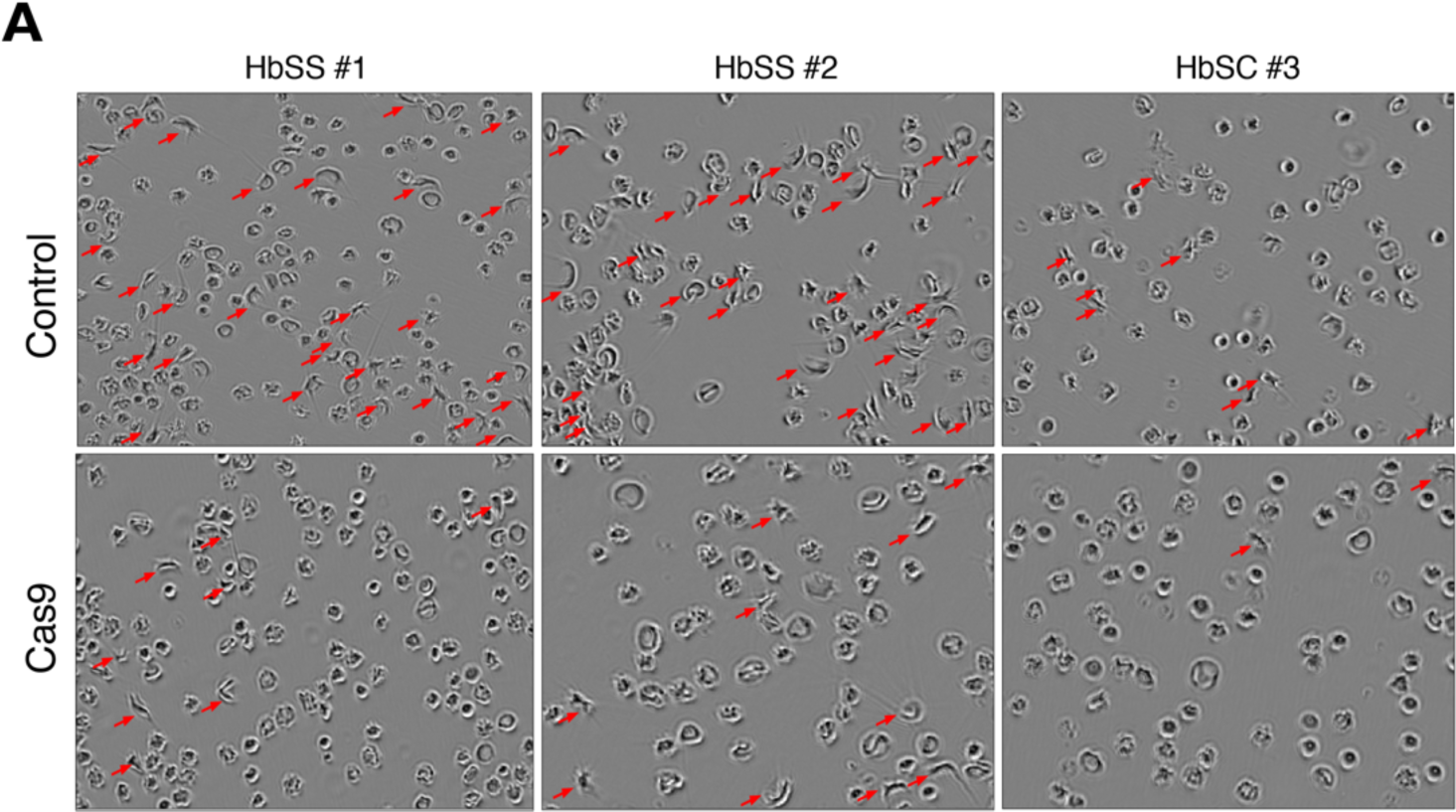
Sickling assay for in vitro differentiated erythroid cells for control and edited human HSPCs derived from three SCD donors. (**A**) In vitro differentiated erythroid cells were sorted with hoechst staining for enucleated cells and incubated for 8 hours in 2% O2 from edited and controls. Images were captured by phase-contrast microscopy using InuCyte S3 Live-cell Analysis System (Sartorius). Sickled cells are indicated with arrows (red).

**Figure S4:**
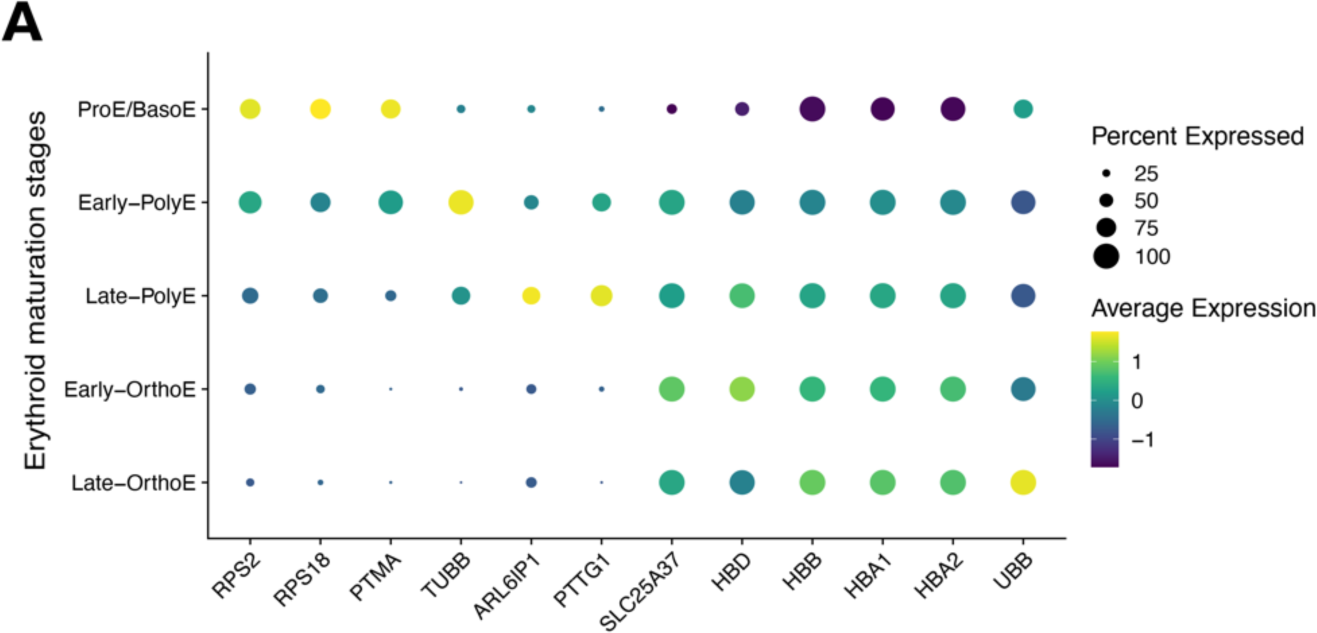
Dot plot showing erythroid maturation markers for scRNA-seq cell type annotation. (**A**) X-axis represents the gene markers and y-axis indicates the terminal stages of erythroid maturation. Dot colors and sizes represent the average gene expression level and the percentages of expressed cells in each cell type.

**Figure S5:**
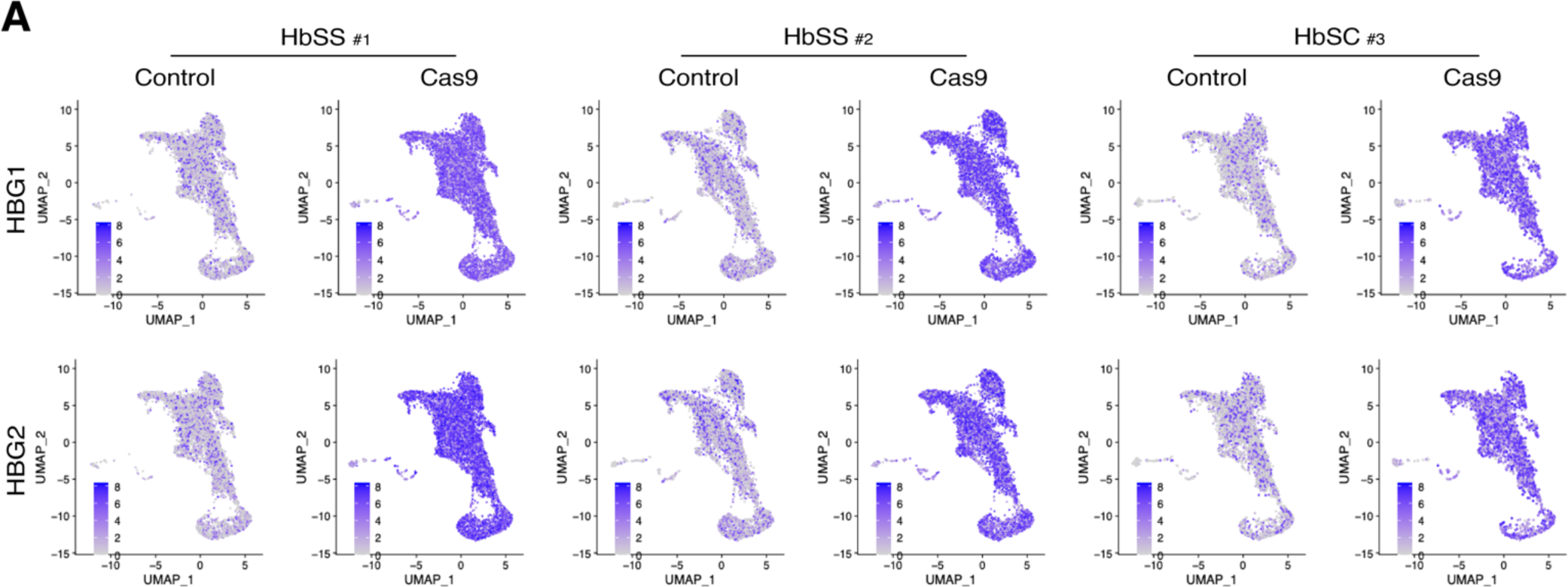
UMAP plots showing HBG1 and HBG2 distributions for individual donors. (A) Y-axis represents the HBG1 and HBG2 transcript-level expression between control and Cas9 edited cells for three SCD donors.

**Figure S6:**
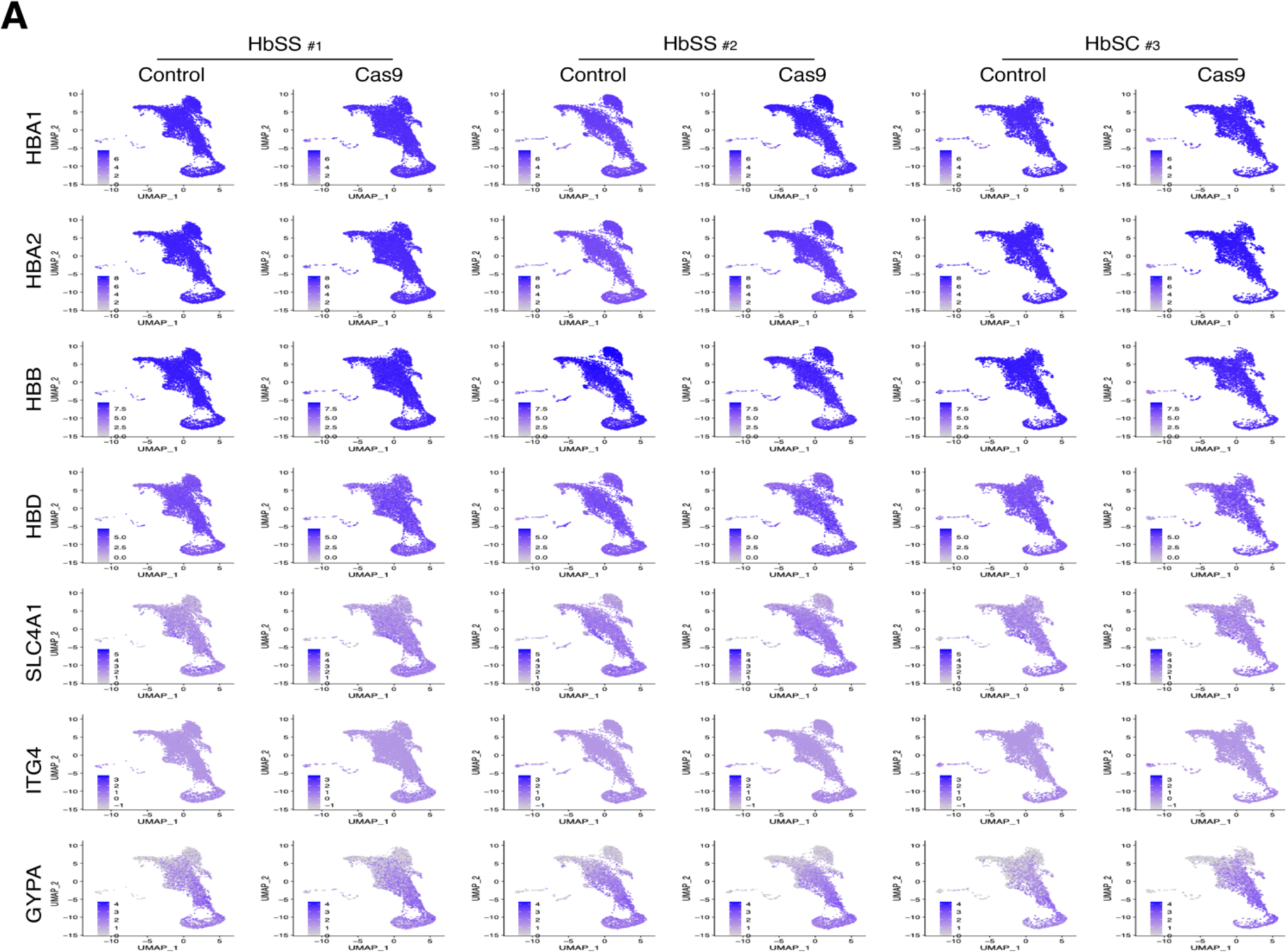
UMAP plots showing marker gene distributions for individual donors. (A) Marker genes of HBA1, HBA2, HBB, HBD, SLC4A1, ITG4, GYPA are represented on rows and three SCD donors of control and Cas9 edited cells are represented on columns.

**Figure S7:**
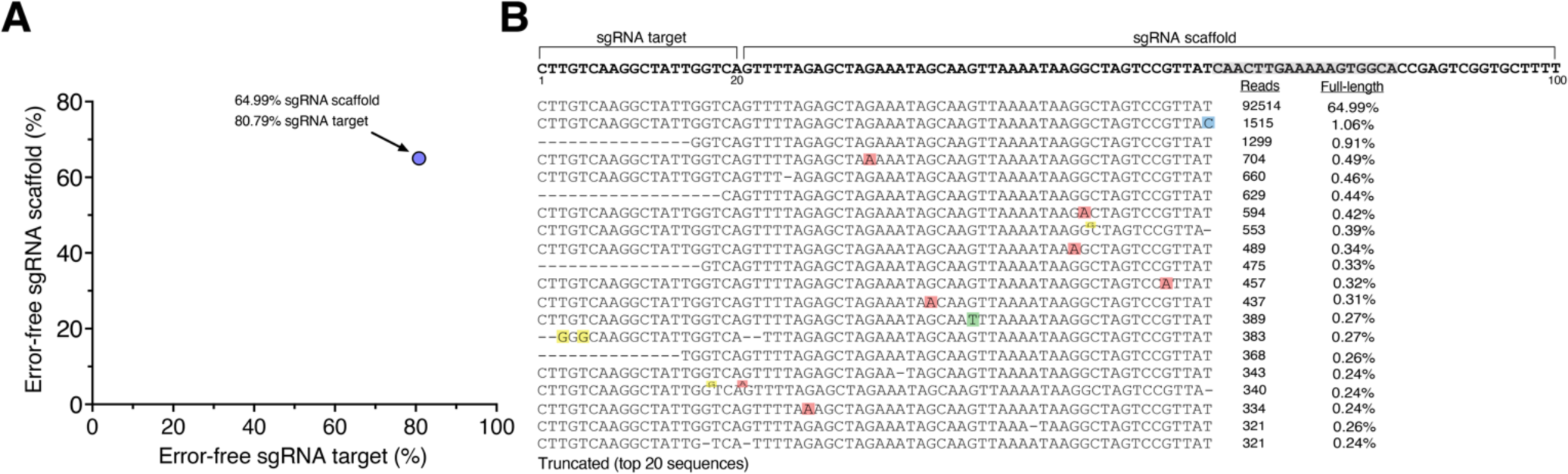
single guide-RNA (sgRNA) sequencing of GMP grade HBG-sgRNA using the smarter smRNA-seq technology. (**A**) Scatter plot shows the percentage of HBG-sgRNA sequence reads mapped to the target sgRNA (shown on x-axis) and full-length sgRNA (shown on y-axis). (**B**) Alignment plot represents the top 20 sequences mapped to the full-length HBG-sgRNA. The first 20bp are the target sgRNA followed by 80bp scaffold sequence. Tailed oligo for first cDNA synthesis binds to the scaffold sgRNA highlighted in orange. Mismatches such as insertions/substitutions are highlighted in blue, red, green, yellow for C, A, T, and G and truncations are highlighted as dashes (−). Columns on right shows the mapped reads to the reference and percentage of perfectly matched sgRNA target and scaffold.

**Figure S8:**
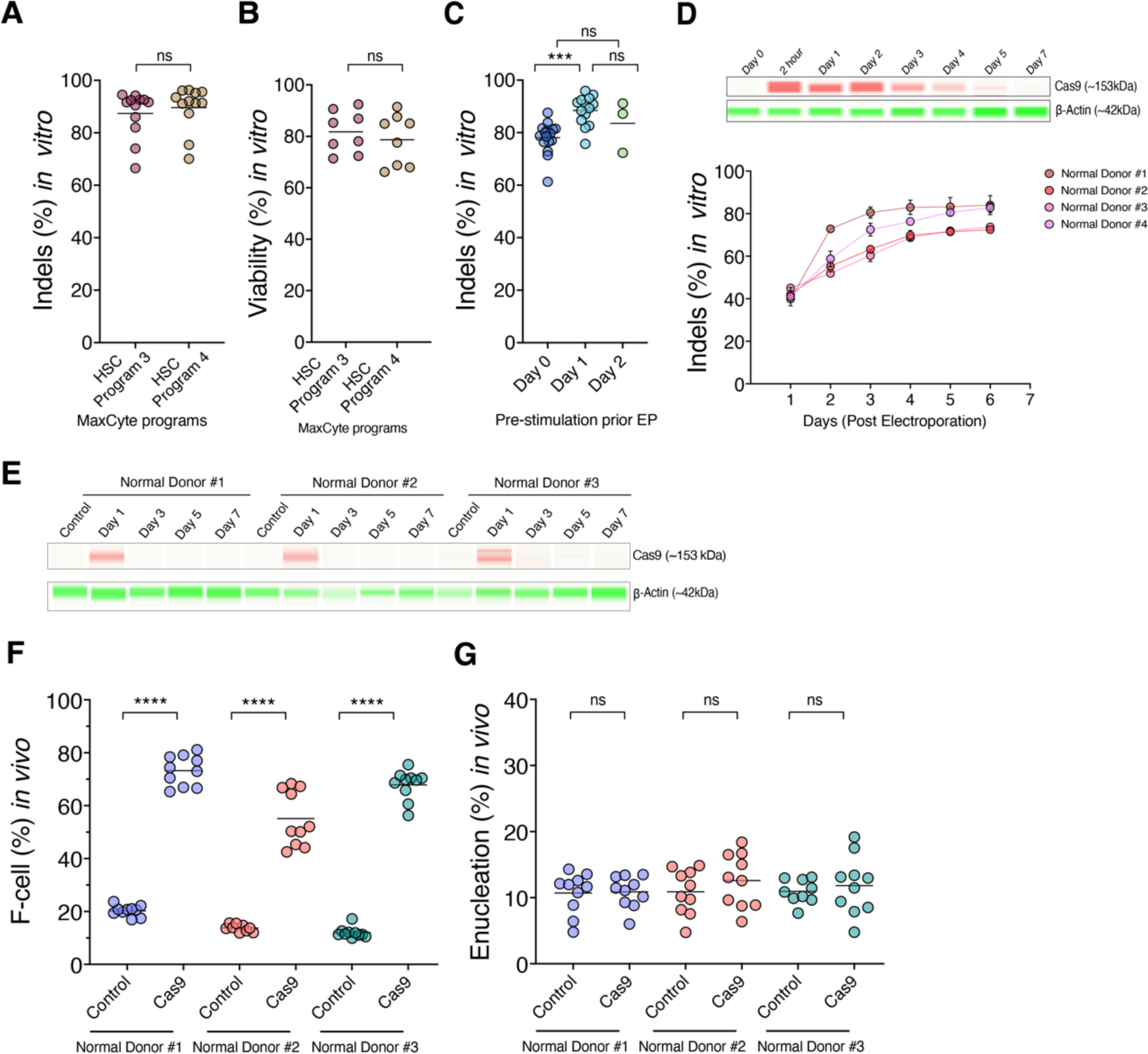
Optimization of −115 g-globin promoter editing in CD34^+^ HSPCs using MaxCyte GTX. (**A**) Editing rate of CD34+ HSPCs with different programs using an OC100 cartridges in the MaxCyte electroporator. (**B**) Viability of edited HSPCs measured after 24 hours of EP. (**C**) Editing rate of freshly isolated, Plerixafor mobilized, CD34+ HSPCs pre-stimulated in cytokine supplemented X-VIVO-10 media for 0-2 days before electroporating in an OC100 Cartridge using the HSC-3 program in the MaxCyte electroporator. (**D**) Persistence of Cas9 in post-electroporated CD34^+^ HSPCs by capillary western blot analysis. Size of Cas9 (red) is ∼153kDa and b-actin (green) is ∼42kDa. Percentages of indels to evaluate editing kinetics after EP (from days 0 to 7). X-axis indicates post-electroporation days and y-axis indicates indels (n=4, GCSF-mobilized normal donors). (**E**) Cas9 persistence in post-thaw of cryopreserved cells after 24 hours of electroporation by capillary western blot analysis from three, normal donors (n=3). (**F)** Measurement of F-cell in human CD235a^+^ fractions derived from mice bone marrow after 17 weeks by flow cytometry (For three donors P values <0.0001 ****). (**G**) Measurement of enucleation in *in vitro* differentiated erythroid cells stained with hoechst staining by flow cytometry. Unpaired t-tests utilized, statistically analysis indicates no significant (ns) changes between control and Cas9 edited cells.

**Figure S9:**
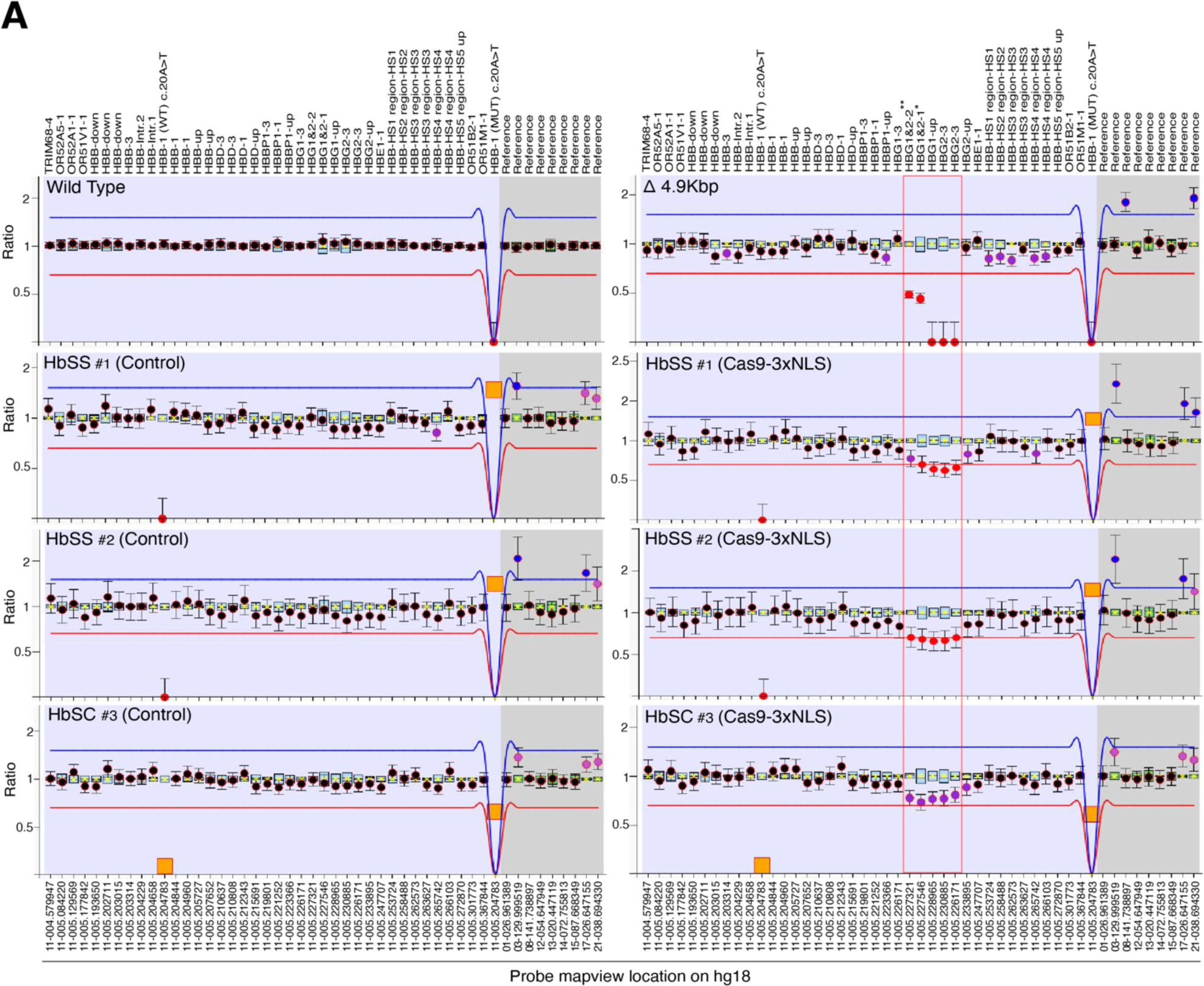
MLPA analysis of bulk bone marrow (BM) at the −115 g-globin gene promoters after 17 weeks transplantation. (**A**) Representation of MLPA ratio charts for human β-globin gene cluster using commercial SALSA-MLPA probe mix P102-D1 HBB. Bulk BM samples derived from three SCD donors at 17 weeks post-transplantation shown as controls (left) and edited as Cas9-3xNLS (right). X-axis represents the probe location on chromosome 11 + distance (nt) from the p-telomere to the start of probe sequence. Corresponding genes were shown at the top panel. Y-axis indicates the dosage quotient (DQ) ratio. Blue and red arbitrary lines represent the normal DQ range between 1.2 and 0.8. Probe HBB-1(Mut) c.20A>T indicate the presence of HbS mutation for homozygous (1.2) or heterozygous (0.8) shown in orange square whereas HBB-1(WT) c.20A>T probe implies no signal in HbSS or HbSC samples. Red dot denotes the possible deletion or mutation and five MLPA probes are highlighted in the red box represents the deletions caused by −115 g-globin gene promoters editing. Wild type and homozygous **Δ**4.9kbp deletion HUDEP2 clones were included as controls. HBG1&2-1 and HBG1&2-2 probes have two binding sites, * probe indicates binding to *HBG1* (5227546-5227612), and *HBG2* (5232468-5232536) and ** probe also targets *HBG1* (5227321-5227386) and *HBG2* (5232245-5232310).

**Figure S10:**
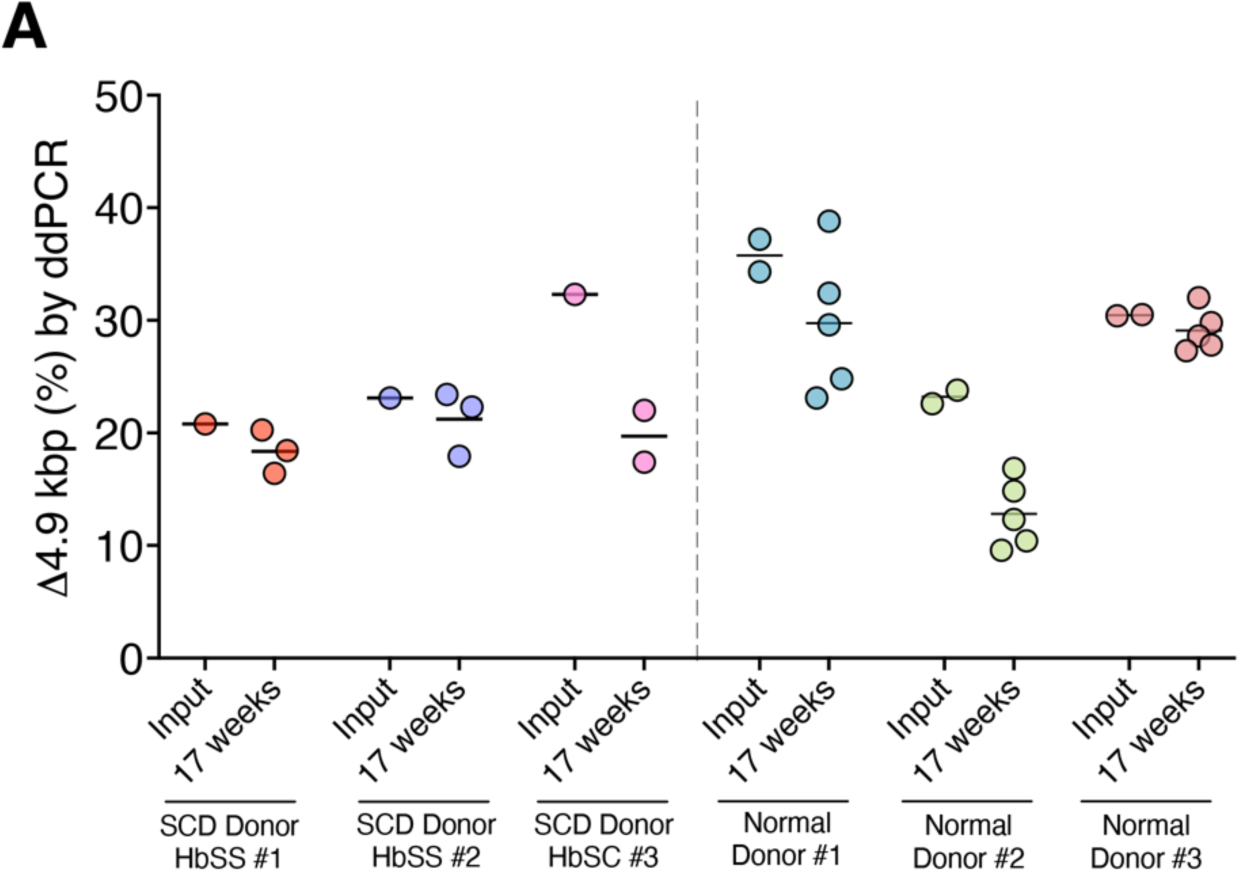
Quantification of 4.9 kb deletion digital droplet PCR (ddPCR) in bulk edited CD34+ HSPCs and bone marrow from mice at 17 weeks. (A) Quantitative measurement of 4.9kbp deletion by digital droplet (ddPCR) in human edited bulk CD34+ HSPCs (input, day 5) and bone marrow derived from mice (17 weeks) in SCD and normal donors.

**Figure S11:**
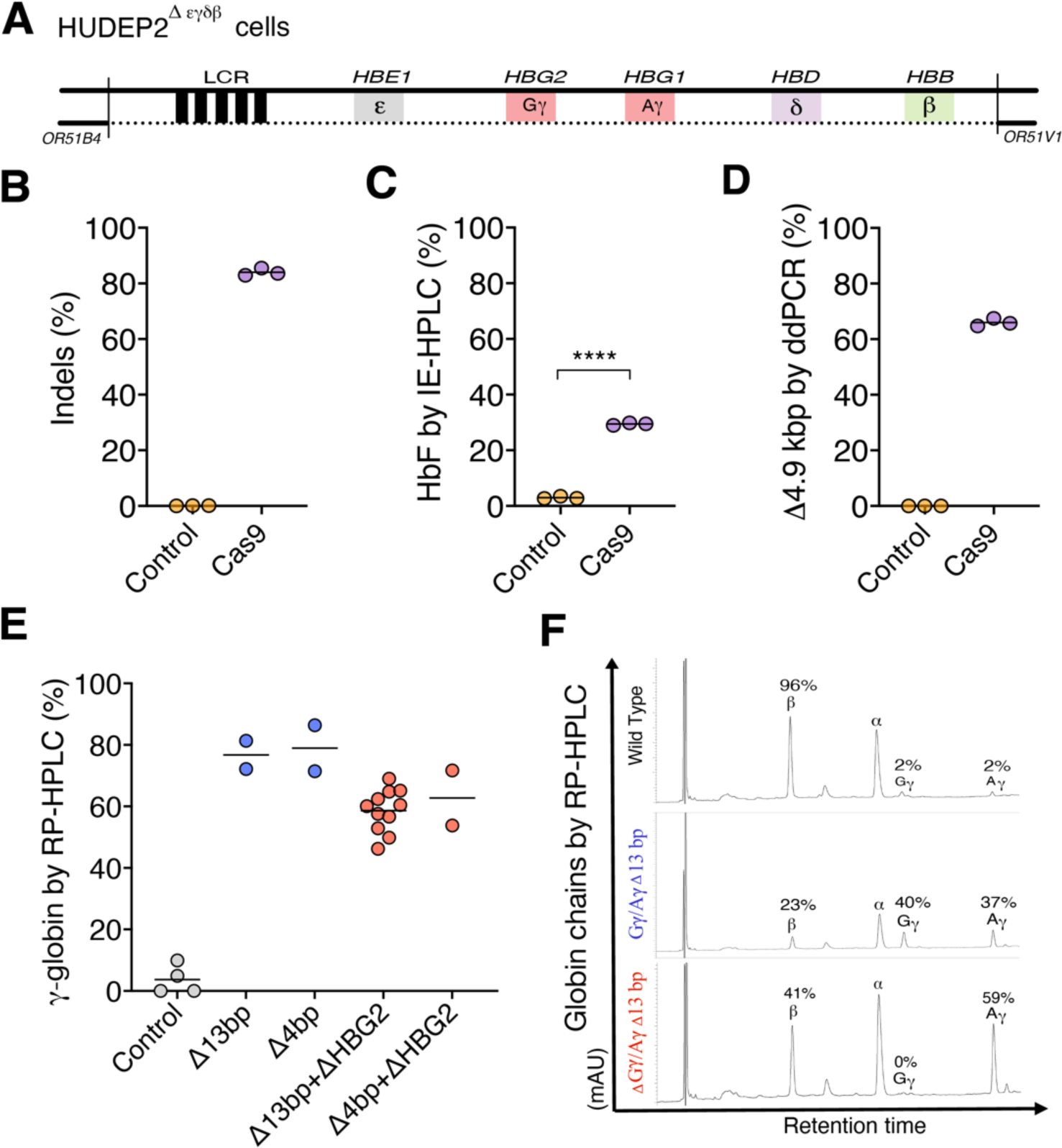
Expression of fetal hemoglobin by editing at the −115 g-globin gene promoters in HUDEP2Degdb clones. (A) Schematic of HBB locus in HUDEP2Degdb cells harboring a heterozygous 91-kb deletion encompassing the extended β-like globin locus were edited with Cas9-3xNLS complexed with HBG-sgRNA. (B) Percentage of indels measured by NGS in bulk edited cells at day 3 post electroporation. (C) Percentage of HbF measured by ion-exchange (IE)-HPLC in erythroid differentiated bulk edited cells at day 10 (Unpaired t-test, P-value <0.0001****. (D) Evaluation of 4.9kb deletion in bulk edited cells by digital droplet PCR (ddPCR). (E) Reverse-phase high performance liquid chromatography (RP-HPLC) chromatograms of lysates from HUDEP2Degdb clones with the indicated genotypes. Dots represents individual clones (blue or red). (F) Representative RP-HPLC chromatograms Gγ and Aγ refer to the protein products of HBG2 and HBG1, respectively.

**Figure S12:**
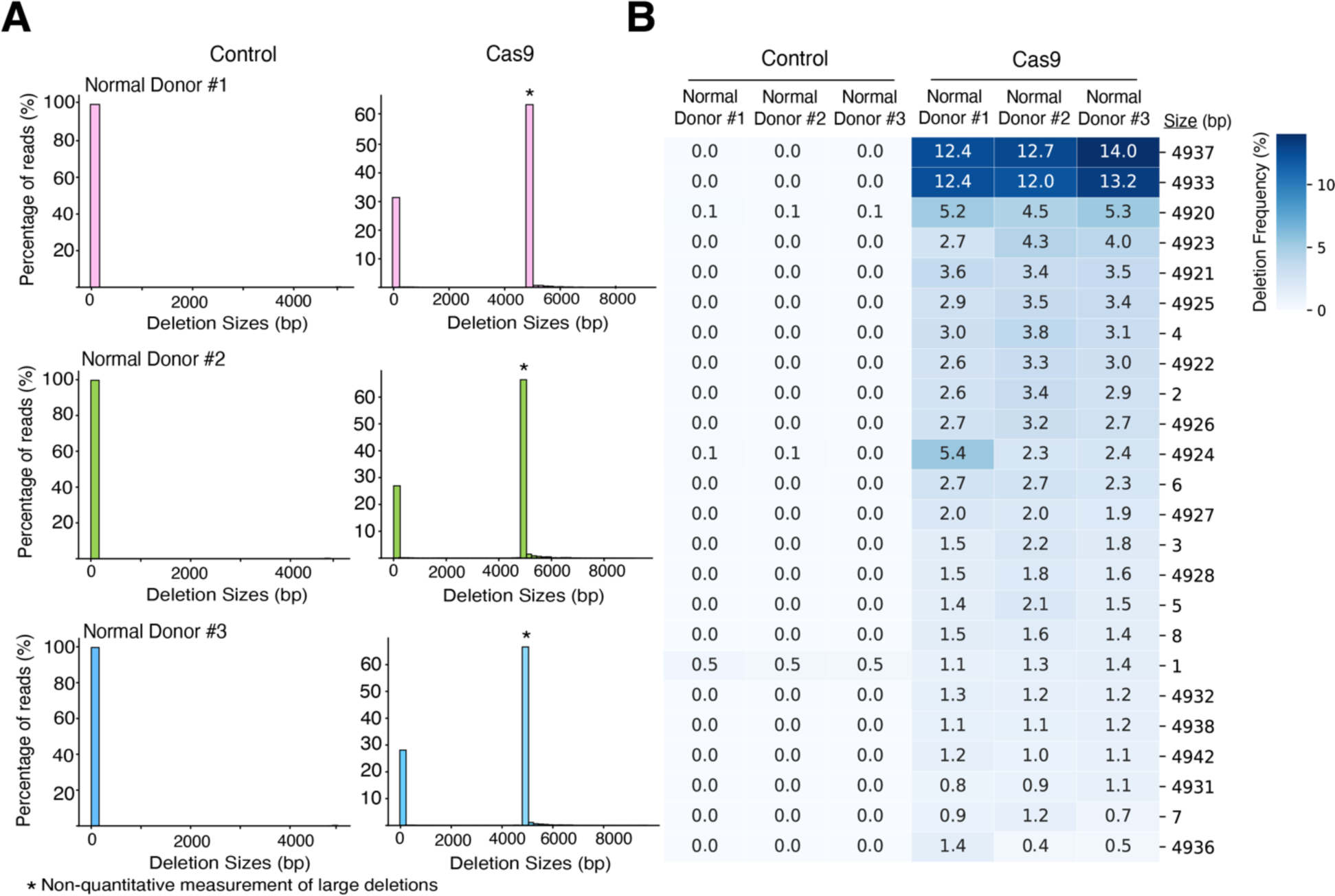
Evaluation of large deletions by long-range PCR based sequencing by PacBio sequencer. (A) Bar plots shows the distribution of deletion size (bp) between control and Cas9 for three normal donors by PacBio sequencing. (B) Heat map shows the frequency of large deletions between control and Cas9. 4.9kbp deletions are more dominant in Cas9 edited CD34+ HSPCs. Note that these frequencies are biased toward smaller PCR products and are non-quantitative.

**Figure S13:**
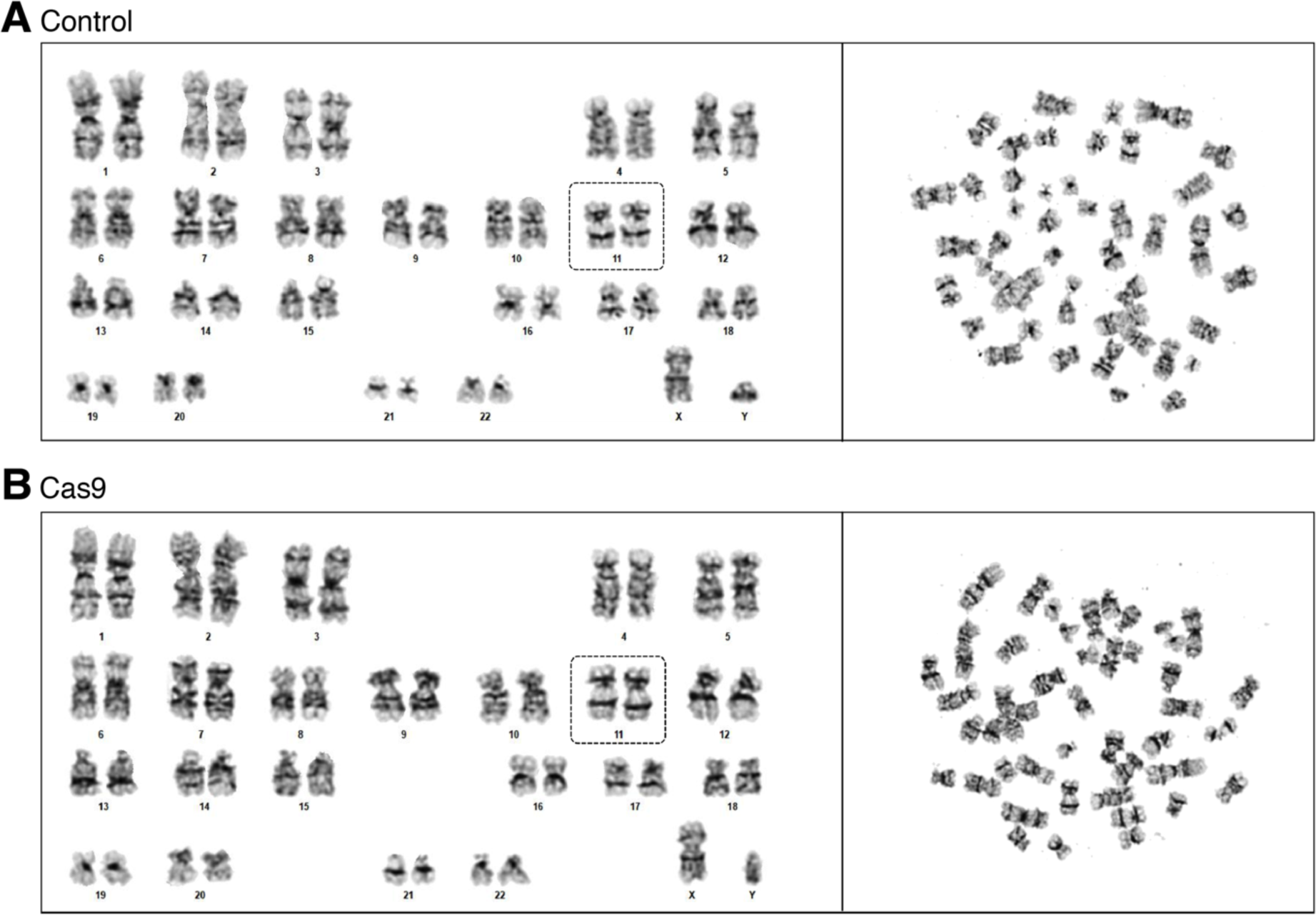
Standard karyotyping of CD34+ HSPCs edited at −115 g-globin gene promoters. Image representative of karyotypic analysis of 20 metaphase cells counted per treatment (n = 4, GCSF mobilized donors). No abnormalities observed with chromosomal morphology and mitotic index between control (A) and Cas9-3xNLS (B). Chromosome 11 is highlighted in dotted box.

**Figure S14:**
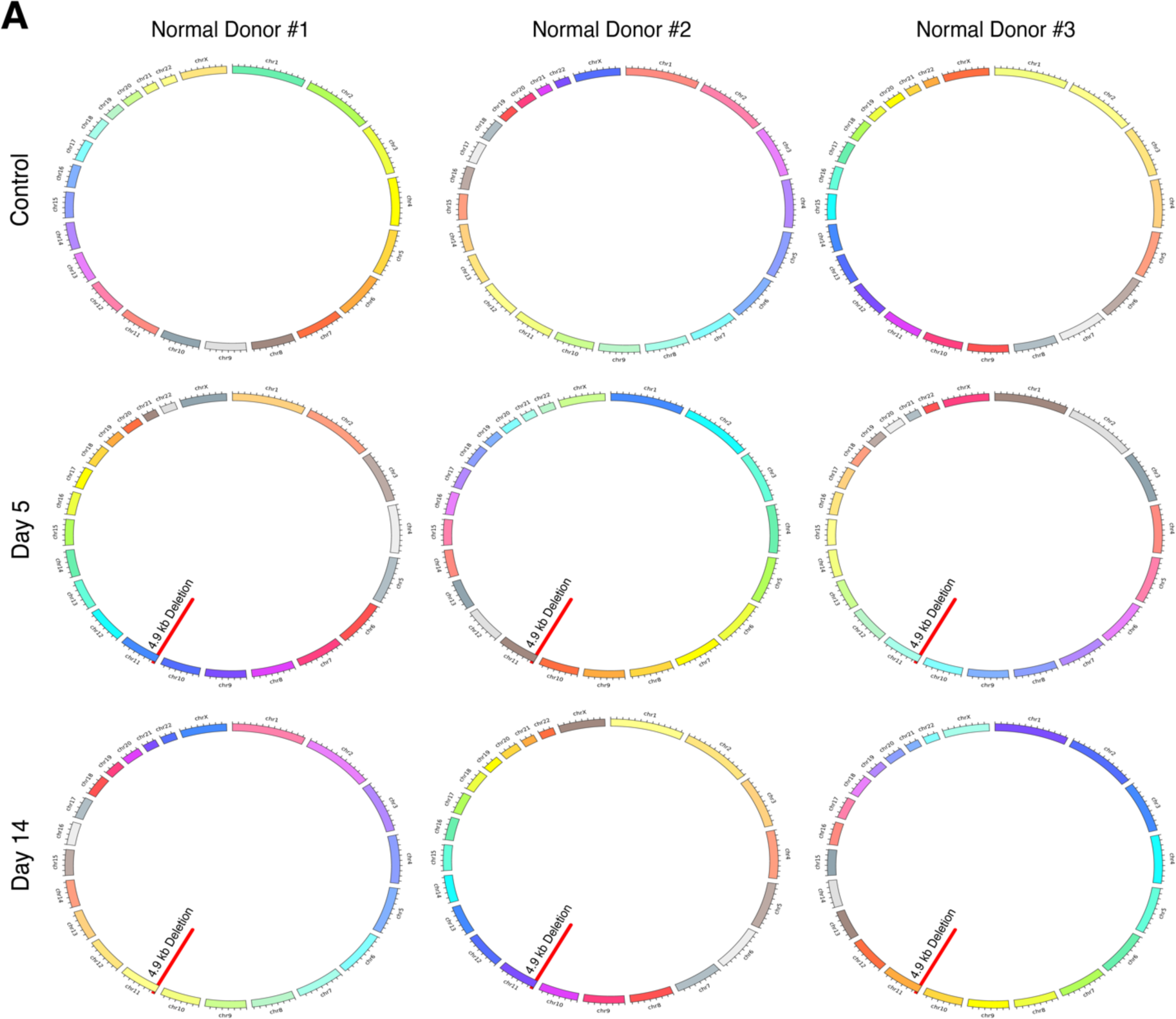
Assessment of genomic re-arrangements in edited CD34^+^ HSPCs at day 5 and day 14 by UDiTaS method. (A) Circos plots representing genomic re-arrangements for −115 g-globin promoter edited CD34+ HSPCs by UDiTaS method (n=3, normal donors). Red line indicates 4.9kbp deletion.

## Tables

**Table S1:**
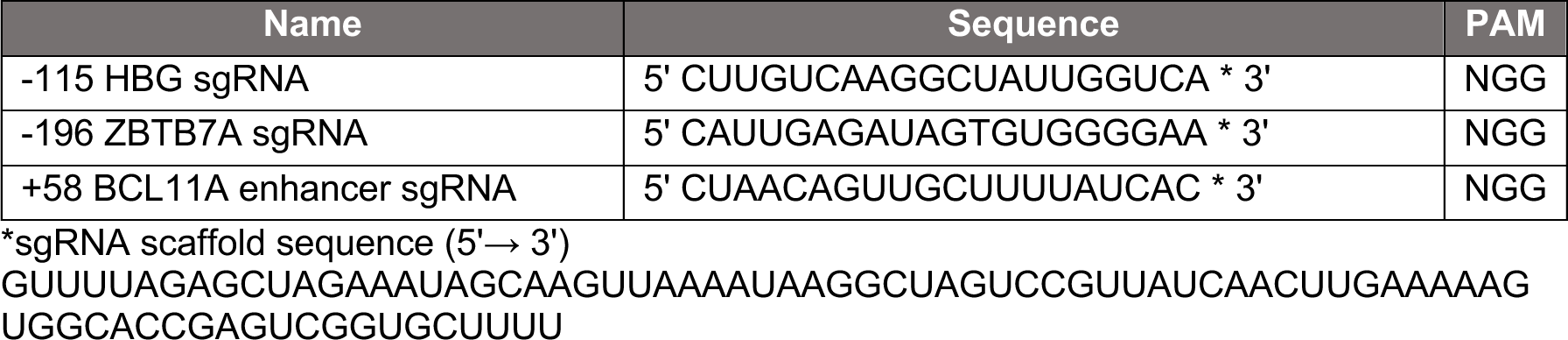
Sequences of chemically modified single guide RNAs (sgRNAs).

**Table S2:**
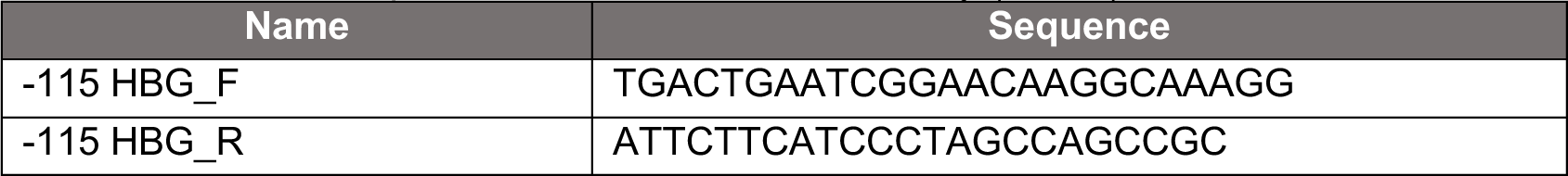
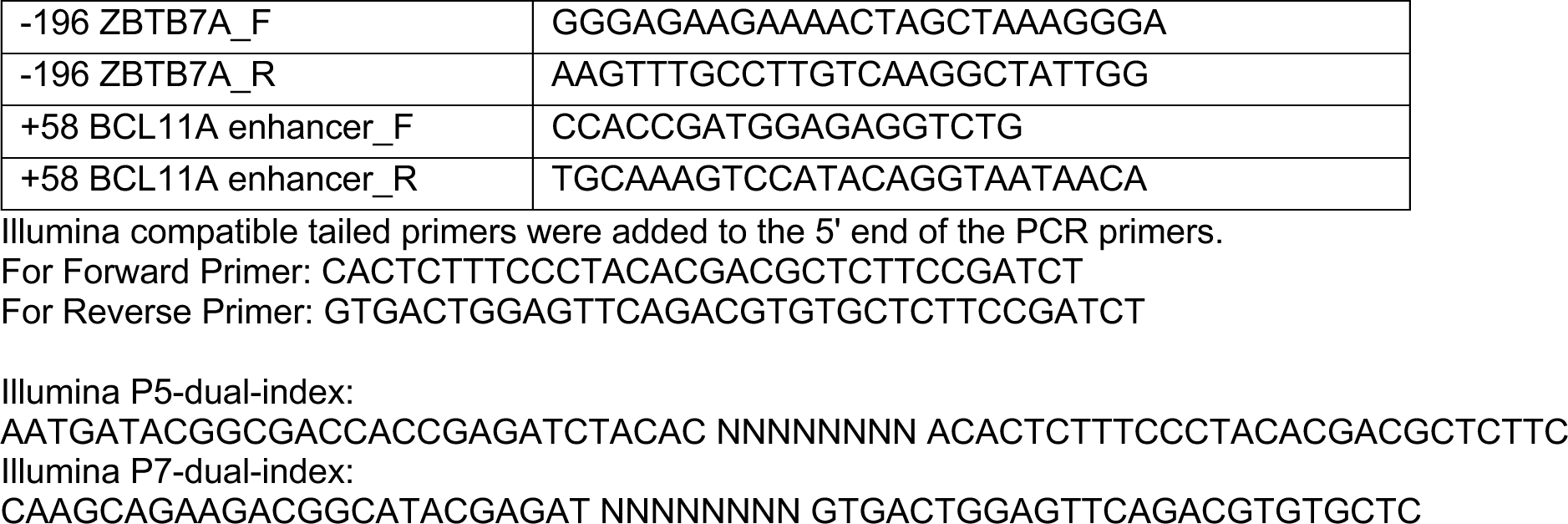
List of PCR primers used for NGS in this study (5’→ 3’).

**Table S3:**
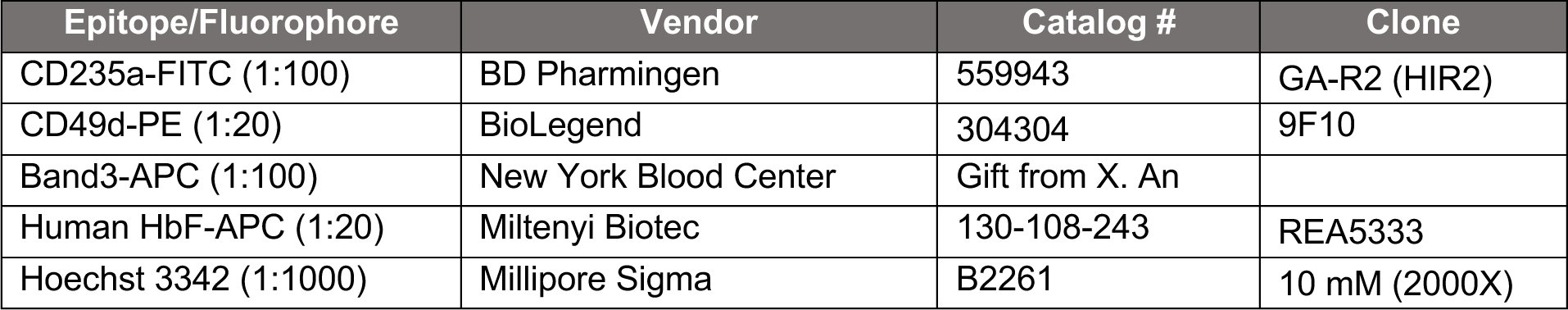
List of antibodies used for flow cytometry for erythroid maturation and enucleation.

**Table S4:**
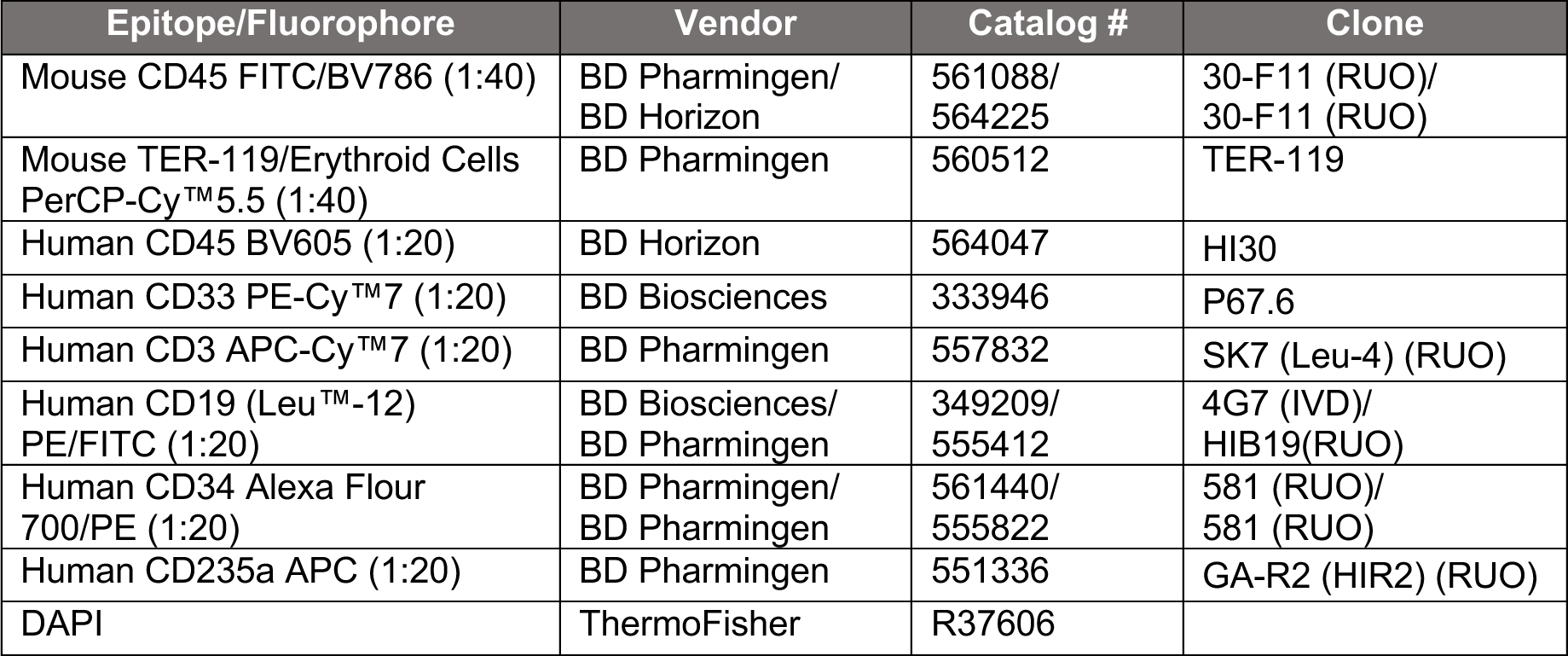
List of antibodies for flow cytometry to measure human engraftment and chimerism in *NBSGW* xenotransplantation.

**Table S5:**
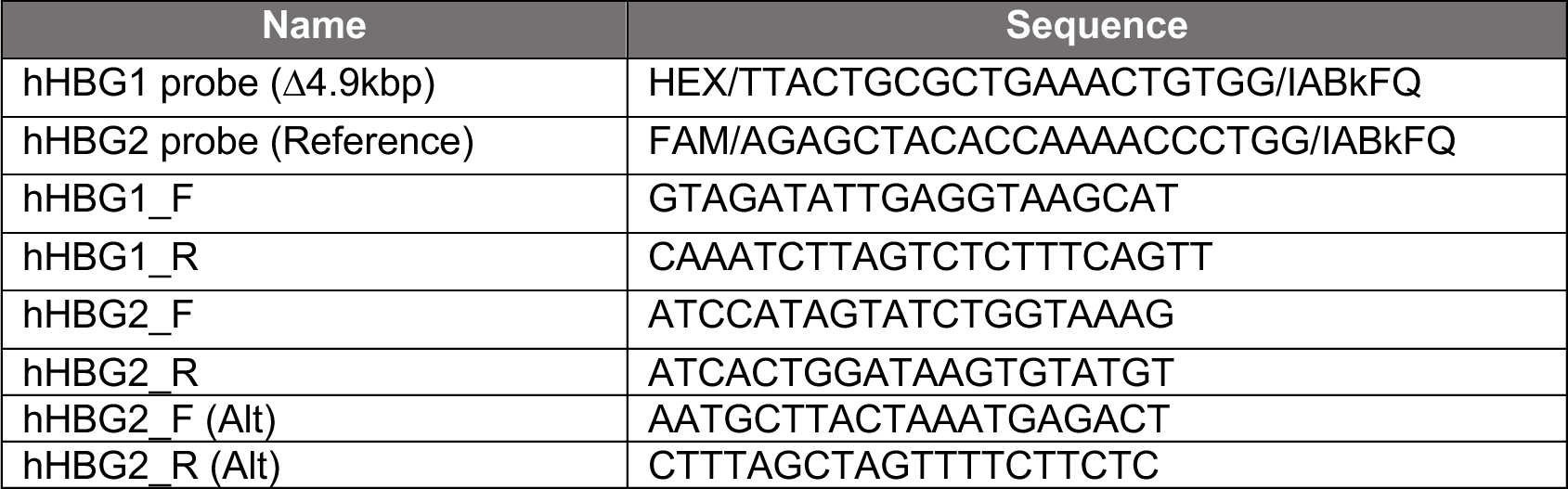
List of digital droplet probes and primers used in this study (5’→ 3’).

**Table S6:**
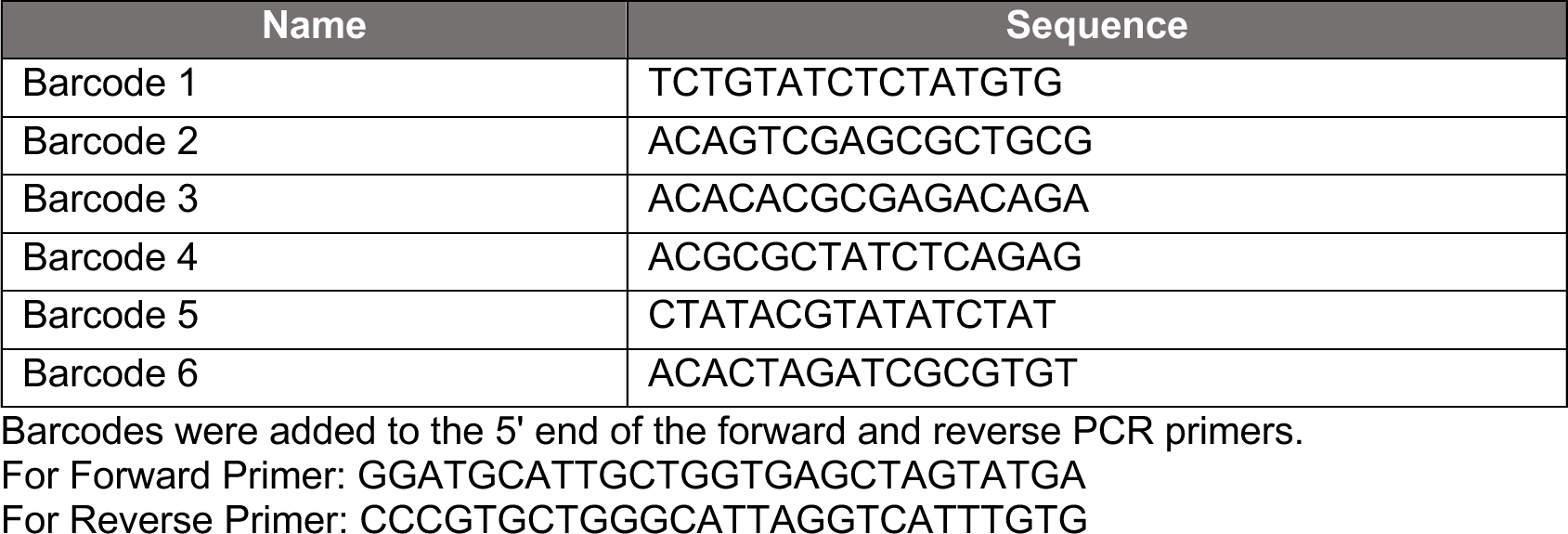
List of PacBio primers with barcodes used in this study (5’→ 3’).

**Table S7:**
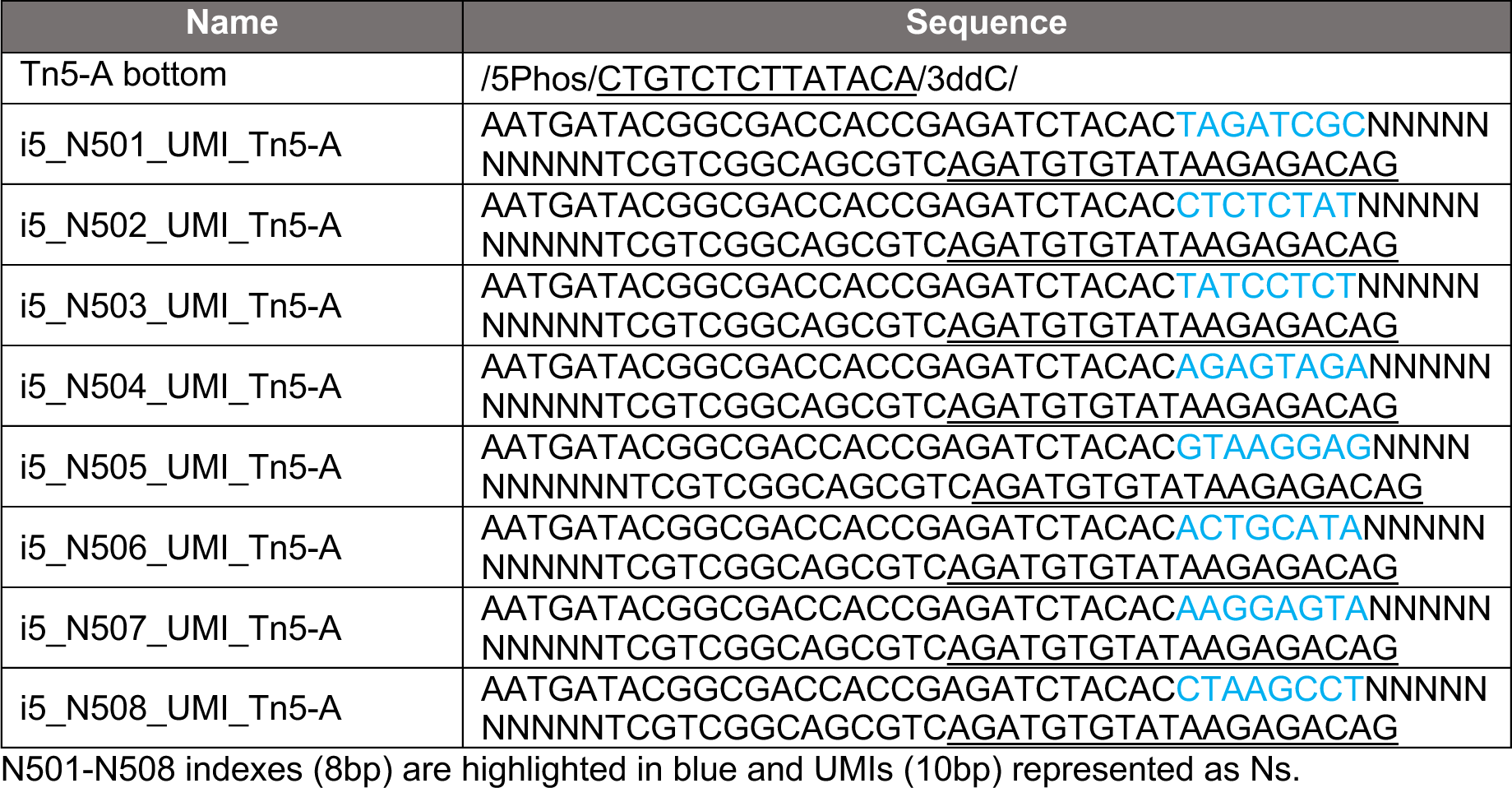
List of Tn5 oligos and primer sequences for UDiTaS protocol.

**Table S8:**
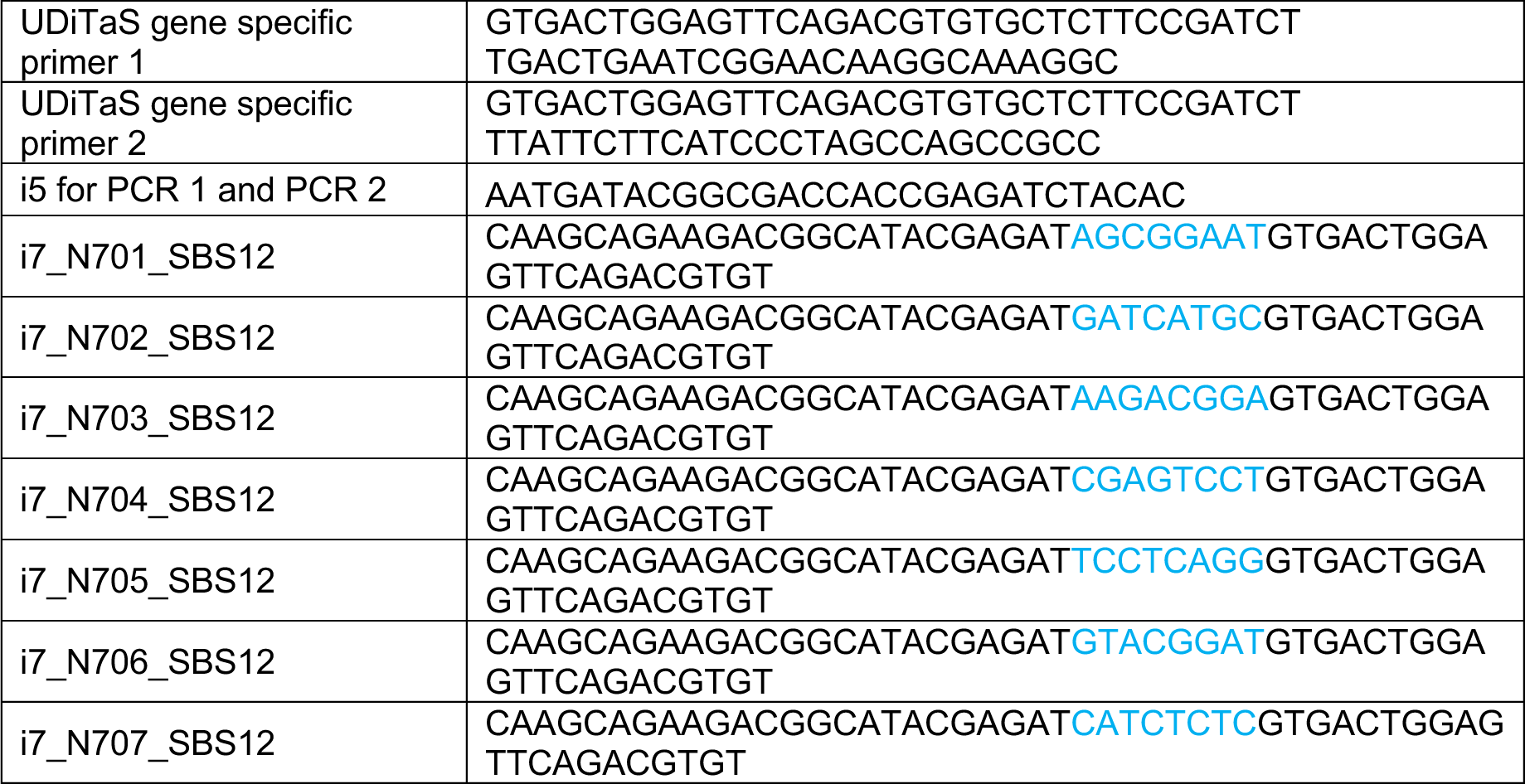
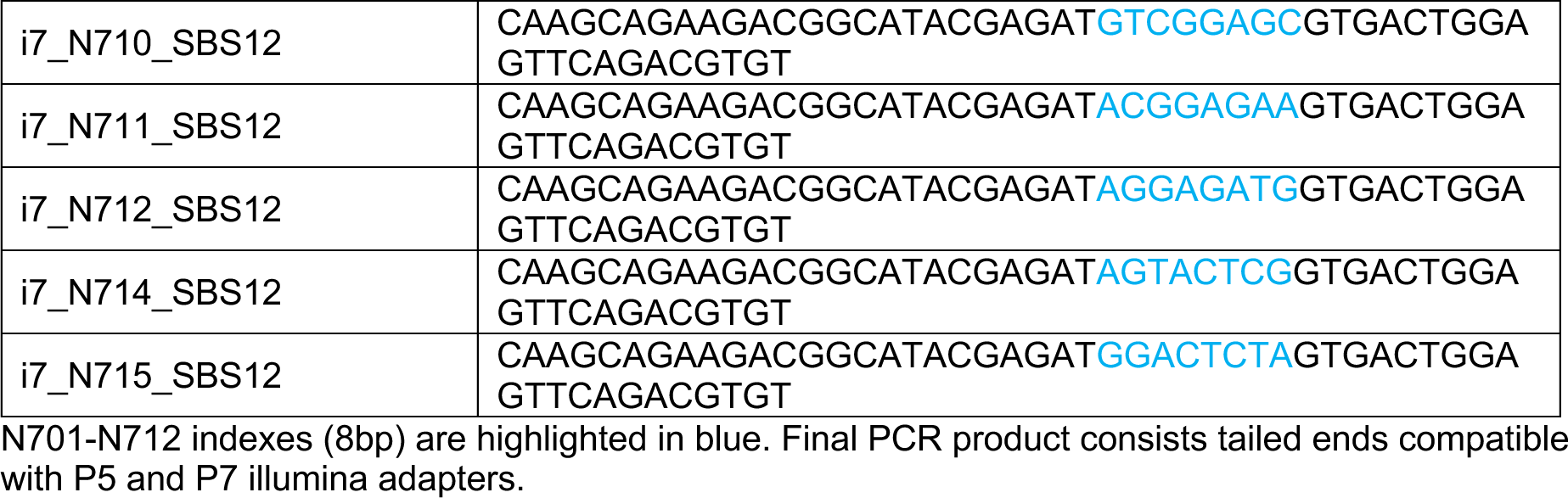
Cas-OFFinder predicted and CHANGE-seq reproducibly identified off-target sites used in rhAmp-seq panel. Excel file.

## References

1. Piel, F.B., Steinberg, M.H., and Rees, D.C. (2017). Sickle cell disease. N. Engl. J. Med. 377, 305.

2. Ingram, V.M. (1956). A specific chemical difference between the globins of normal human and sickle-cell anaemia haemoglobin. Nature 178, 792–794.

3. Bunn, H.F. (1997). Pathogenesis and treatment of sickle cell disease. N. Engl. J. Med. 337, 762–769.

4. Kato, G.J., Piel, F.B., Reid, C.D., Gaston, M.H., Ohene-Frempong, K., Krishnamurti, L., Smith, W.R., Panepinto, J.A., Weatherall, D.J., Costa, F.F., et al. (2018). Sickle cell disease. Nat. Rev. Dis. Primers 4, 18011.

5. Migotsky, M., Beestrum, M., and Badawy, S.M. (2022). Recent advances in sickle-cell disease therapies: A review of voxelotor, crizanlizumab, and L-glutamine. Pharmacy (Basel) 10. 10.3390/pharmacy10050123.

6. Walters, M.C., Patience, M., Leisenring, W., Eckman, J.R., Buchanan, G.R., Rogers, Z.R., Olivieri, N.E., Vichinsky, E., Davies, S.C., Mentzer, W.C., et al. (1996). Barriers to bone marrow transplantation for sickle cell anemia. Biol. Blood Marrow Transplant. 2, 100–104.

7. Eapen, M., Brazauskas, R., Walters, M.C., Bernaudin, F., Bo-Subait, K., Fitzhugh, C.D., Hankins, J.S., Kanter, J., Meerpohl, J.J., Bolaños-Meade, J., et al. (2019). Effect of donor type and conditioning regimen intensity on allogeneic transplantation outcomes in patients with sickle cell disease: a retrospective multicentre, cohort study. Lancet Haematol. 6, e585–e596.

8. Doerfler, P.A., Sharma, A., Porter, J.S., Zheng, Y., Tisdale, J.F., and Weiss, M.J. (2021). Genetic therapies for the first molecular disease. J. Clin. Invest. 131. 10.1172/JCI146394.

9. Kanter, J., Thompson, A.A., Pierciey, F.J., Jr, Hsieh, M., Uchida, N., Leboulch, P., Schmidt, M., Bonner, M., Guo, R., Miller, A., et al. (2023). Lovo-cel gene therapy for sickle cell disease: Treatment process evolution and outcomes in the initial groups of the HGB-206 study. Am. J. Hematol. 98, 11–22.

10. Frangoul, H., Ho, T.W., and Corbacioglu, S. (2021). CRISPR-Cas9 gene editing for sickle cell disease and β-thalassemia. Reply. N. Engl. J. Med. 384, e91.

11. Wilkinson, A.C., Dever, D.P., Baik, R., Camarena, J., Hsu, I., Charlesworth, C.T., Morita, C., Nakauchi, H., and Porteus, M.H. (2021). Cas9-AAV6 gene correction of beta-globin in autologous HSCs improves sickle cell disease erythropoiesis in mice. Nat. Commun. 12, 686.

12. Newby, G.A., Yen, J.S., Woodard, K.J., Mayuranathan, T., Lazzarotto, C.R., Li, Y., Sheppard-Tillman, H., Porter, S.N., Yao, Y., Mayberry, K., et al. (2021). Base editing of haematopoietic stem cells rescues sickle cell disease in mice. Nature 595, 295–302.

13. Frangoul, H., Locatelli, F., Bhatia, M., Mapara, M.Y., Molinari, L., Sharma, A., Lobitz, S., de Montalembert, M., Rondelli, D., Steinberg, M., et al. (2022). Efficacy and safety of a single dose of exagamglogene autotemcel for severe sickle cell disease. Blood 140, 29–31.

14. Leonard, A., and Tisdale, J.F. (2021). A pause in gene therapy: Reflecting on the unique challenges of sickle cell disease. Mol. Ther. 29, 1355–1356.

15. Magrin, E., Miccio, A., and Cavazzana, M. (2019). Lentiviral and genome-editing strategies for the treatment of β-hemoglobinopathies. Blood 134, 1203–1213.

16. Frati, G., and Miccio, A. (2021). Genome editing for β-hemoglobinopathies: Advances and challenges. J. Clin. Med. 10, 482.

17. Romero, Z., Lomova, A., Said, S., Miggelbrink, A., Kuo, C.Y., Campo-Fernandez, B., Hoban, M.D., Masiuk, K.E., Clark, D.N., Long, J., et al. (2019). Editing the sickle cell disease mutation in human hematopoietic stem cells: Comparison of endonucleases and homologous donor templates. Mol. Ther. 27, 1389–1406.

18. Ferrari, S., Jacob, A., Cesana, D., Laugel, M., Beretta, S., Varesi, A., Unali, G., Conti, A., Canarutto, D., Albano, L., et al. (2022). Choice of template delivery mitigates the genotoxic risk and adverse impact of editing in human hematopoietic stem cells. Cell Stem Cell 29, 1428–1444.e9.

19. Fiumara, M., Ferrari, S., Omer-Javed, A., Beretta, S., Albano, L., Canarutto, D., Varesi, A., Gaddoni, C., Brombin, C., Cugnata, F., et al. (2023). Genotoxic effects of base and prime editing in human hematopoietic stem cells. Nat. Biotechnol. 10.1038/s41587-023-01915-4.

20. Steinberg, M.H. (2020). Fetal hemoglobin in sickle cell anemia. Blood 136, 2392–2400.

21. Sankaran, V.G., Xu, J., and Orkin, S.H. (2010). Advances in the understanding of haemoglobin switching. Br. J. Haematol. 149, 181–194.

22. Lettre, G., and Bauer, D.E. (2016). Fetal haemoglobin in sickle-cell disease: from genetic epidemiology to new therapeutic strategies. Lancet 387, 2554–2564.

23. Eaton, W.A., and Bunn, H.F. (2017). Treating sickle cell disease by targeting HbS polymerization. Blood 129, 2719–2726.

24. Jacob, G.F., and Raper, A.B. (1958). Hereditary persistence of foetal haemoglobin production, and its interaction with the sickle-cell trait. Br. J. Haematol. 4, 138–149.

25. Forget, B.G. (1998). Molecular basis of hereditary persistence of fetal hemoglobin. Ann. N. Y. Acad. Sci. 850, 38–44.

26. Crossley, M., Christakopoulos, G.E., and Weiss, M.J. (2022). Effective therapies for sickle cell disease: are we there yet? Trends Genet. 38, 1284–1298.

27. Traxler, E.A., Yao, Y., Wang, Y.-D., Woodard, K.J., Kurita, R., Nakamura, Y., Hughes, J.R., Hardison, R.C., Blobel, G.A., Li, C., et al. (2016). A genome-editing strategy to treat β-hemoglobinopathies that recapitulates a mutation associated with a benign genetic condition. Nat. Med. 22, 987–990.

28. Sankaran, V.G., Menne, T.F., Xu, J., Akie, T.E., Lettre, G., Van Handel, B., Mikkola, H.K.A., Hirschhorn, J.N., Cantor, A.B., and Orkin, S.H. (2008). Human fetal hemoglobin expression is regulated by the developmental stage-specific repressor BCL11A. Science 322, 1839–1842.

29. Masuda, T., Wang, X., Maeda, M., Canver, M.C., Sher, F., Funnell, A.P.W., Fisher, C., Suciu, M., Martyn, G.E., Norton, L.J., et al. (2016). Transcription factors LRF and BCL11A independently repress expression of fetal hemoglobin. Science 351, 285–289.

30. Canver, M.C., Smith, E.C., Sher, F., Pinello, L., Sanjana, N.E., Shalem, O., Chen, D.D., Schupp, P.G., Vinjamur, D.S., Garcia, S.P., et al. (2015). BCL11A enhancer dissection by Cas9-mediated in situ saturating mutagenesis. Nature 527, 192–197.

31. Soni, S., Frangoul, H., Bobruff, Y., Cappellini, M.D., Corbacioglu, S., Fernandez, C.M., de la Fuente, J., Grupp, S., Handgretinger, R., Ho, T.W., et al. (2021). Safety and efficacy of CTX001 in patients with transfusion-dependent β-thalassemia (TDT) or sickle cell disease (SCD): Early results from the climb THAL-111 and climb SCD-121 studies of autologous CRISPR-Cas9-modified CD34+ hematopoietic stem and progenitor cells (HSPCs). Transplantation and Cellular Therapy 27, S72–S73.

32. Fu, B., Liao, J., Chen, S., Li, W., Wang, Q., Hu, J., Yang, F., Hsiao, S., Jiang, Y., Wang, L., et al. (2022). CRISPR-Cas9-mediated gene editing of the BCL11A enhancer for pediatric β0/β0 transfusion-dependent β-thalassemia. Nat. Med. 28, 1573–1580.

33. Métais, J.-Y., Doerfler, P.A., Mayuranathan, T., Bauer, D.E., Fowler, S.C., Hsieh, M.M., Katta, V., Keriwala, S., Lazzarotto, C.R., Luk, K., et al. (2019). Genome editing of HBG1 and HBG2 to induce fetal hemoglobin. Blood Adv 3, 3379–3392.

34. Chang, K.-H., Sanchez, M., Heath, J., deDreuzy, E., Haskett, S., Vogelaar, A., Gogi, K., Da Silva, J., Wang, T., Sadowski, A., et al. (2018). Comparative studies reveal robust HbF induction by editing of HBG1/2 promoters or BCL11A erythroid-enhancer in human CD34+ cells but that BCL11A erythroid-enhancer editing is associated with selective reduction in erythroid lineage reconstitution in a xenotransplantation model. Blood 132, 409–409.

35. Heath, J., de Dreuzy, E., Sanchez, M., Haskett, S., Wang, T., Sousa, P., Gotta, G., Siwak, J., Viswanathan, R., Ta, F., et al. (2019). Ps1518 genome editing of hbg1/2 promoter leads to robust hbf induction in vivo, while editing of bcl11a erythroid enhancer results in erythroid defects. HemaSphere 3, 699–700.

36. Wienert, B., Martyn, G.E., Funnell, A.P.W., Quinlan, K.G.R., and Crossley, M. (2018). Wake-up Sleepy Gene: Reactivating Fetal Globin for β-Hemoglobinopathies. Trends Genet. 34, 927–940.

37. Vakulskas, C.A., Dever, D.P., Rettig, G.R., Turk, R., Jacobi, A.M., Collingwood, M.A., Bode, N.M., McNeill, M.S., Yan, S., Camarena, J., et al. (2018). A high-fidelity Cas9 mutant delivered as a ribonucleoprotein complex enables efficient gene editing in human hematopoietic stem and progenitor cells. Nat. Med. 24, 1216–1224.

38. Wu, Y., Zeng, J., Roscoe, B.P., Liu, P., Yao, Q., Lazzarotto, C.R., Clement, K., Cole, M.A., Luk, K., Baricordi, C., et al. (2019). Highly efficient therapeutic gene editing of human hematopoietic stem cells. Nat. Med. 25, 776–783.

39. Hu, Z., Van Rooijen, N., and Yang, Y.-G. (2011). Macrophages prevent human red blood cell reconstitution in immunodeficient mice. Blood 118, 5938–5946.

40. Powars, D.R., Weiss, J.N., Chan, L.S., and Schroeder, W.A. (1984). Is there a threshold level of fetal hemoglobin that ameliorates morbidity in sickle cell anemia? Blood 63, 921–926.

41. Steinberg, M.H., Chui, D.H.K., Dover, G.J., Sebastiani, P., and Alsultan, A. (2014). Fetal hemoglobin in sickle cell anemia: a glass half full? Blood 123, 481–485.

42. Akinsheye, I., Alsultan, A., Solovieff, N., Ngo, D., Baldwin, C.T., Sebastiani, P., Chui, D.H.K., and Steinberg, M.H. (2011). Fetal hemoglobin in sickle cell anemia. Blood 118, 19–27.

43. Peri, K.G., Gagnon, C., and Bard, H. (1998). Quantitative correlation between globin mRNAs and synthesis of fetal and adult hemoglobins during hemoglobin switchover in the perinatal period. Pediatr. Res. 43, 504–508.

44. Stuart, T., Butler, A., Hoffman, P., Hafemeister, C., Papalexi, E., Mauck, W.M., 3rd, Hao, Y., Stoeckius, M., Smibert, P., and Satija, R. (2019). Comprehensive integration of single-cell data. Cell 177, 1888–1902.e21.

45. Huang, P., Zhao, Y., Zhong, J., Zhang, X., Liu, Q., Qiu, X., Chen, S., Yan, H., Hillyer, C., Mohandas, N., et al. (2020). Putative regulators for the continuum of erythroid differentiation revealed by single-cell transcriptome of human BM and UCB cells. Proc. Natl. Acad. Sci. U. S. A. 117, 12868–12876.

46. Steinberg, M.H., Forget, B.G., Higgs, D.R., and Weatherall, D.J. (2009). Disorders of Hemoglobin: Genetics, Pathophysiology, Clinical Management (United Kingdom Cambridge University Press).

47. Jinek, M., Chylinski, K., Fonfara, I., Hauer, M., Doudna, J.A., and Charpentier, E. (2012). A programmable dual-RNA-guided DNA endonuclease in adaptive bacterial immunity. Science 337, 816–821.

48. Hendel, A., Bak, R.O., Clark, J.T., Kennedy, A.B., Ryan, D.E., Roy, S., Steinfeld, I., Lunstad, B.D., Kaiser, R.J., Wilkens, A.B., et al. (2015). Chemically modified guide RNAs enhance CRISPR-Cas genome editing in human primary cells. Nat. Biotechnol. 33, 985–989.

49. Macias, L.A., Garcia, S.P., Back, K.M., Wu, Y., Johnson, G.H., Kathiresan, S., Bellinger, A.M., Rohde, E., Freitas, M.A., and Madsen, J.A. (2023). Spacer fidelity assessments of guide RNA by top-down mass spectrometry. ACS Cent. Sci. 9, 1437–1452.

50. Mehta, A., and Merkel, O.M. (2020). Immunogenicity of Cas9 Protein. J. Pharm. Sci. 109, 62–67.

51. Center for Biologics Evaluation, and Research Human Gene Therapy Products Incorporating Human Genome Editing. U.S. Food and Drug Administration. https://www.fda.gov/regulatory-information/search-fda-guidance-documents/human-gene-therapy-products-incorporating-human-genome-editing.

52. Doerfler, P.A., Feng, R., Li, Y., Palmer, L.E., Porter, S.N., Bell, H.W., Crossley, M., Pruett-Miller, S.M., Cheng, Y., and Weiss, M.J. (2021). Activation of γ-globin gene expression by GATA1 and NF-Y in hereditary persistence of fetal hemoglobin. Nat. Genet. 53, 1177–1186.

53. Mayuranathan, T., Newby, G.A., Feng, R., Yao, Y., Mayberry, K.D., Lazzarotto, C.R., Li, Y., Levine, R.M., Nimmagadda, N., Dempsey, E., et al. (2023). Potent and uniform fetal hemoglobin induction via base editing. Nat. Genet. 55, 1210–1220.

54. Lazzarotto, C.R., Malinin, N.L., Li, Y., Zhang, R., Yang, Y., Lee, G., Cowley, E., He, Y., Lan, X., Jividen, K., et al. (2020). CHANGE-seq reveals genetic and epigenetic effects on CRISPR–Cas9 genome-wide activity. Nat. Biotechnol. 38, 1317–1327.

55. Giannoukos, G., Ciulla, D.M., Marco, E., Abdulkerim, H.S., Barrera, L.A., Bothmer, A., Dhanapal, V., Gloskowski, S.W., Jayaram, H., Maeder, M.L., et al. (2018). UDiTaS^TM^, a genome editing detection method for indels and genome rearrangements. BMC Genomics 19, 212.

56. Sharma, A., Boelens, J.-J., Cancio, M., Hankins, J.S., Bhad, P., Azizy, M., Lewandowski, A., Zhao, X., Chitnis, S., Peddinti, R., et al. (2023). CRISPR-Cas9 editing of the HBG1 and HBG2 promoters to treat sickle cell disease. N. Engl. J. Med. 389, 820–832.

57. Uchida, N., Leonard, A., Stroncek, D., Panch, S.R., West, K., Molloy, E., Hughes, T.E., Hauffe, S., Taylor, T., Fitzhugh, C., et al. (2020). Safe and efficient peripheral blood stem cell collection in patients with sickle cell disease using plerixafor. Haematologica 105, e497.

58. Sharma, A., Leonard, A., West, K., Gossett, J.M., Uchida, N., Panch, S., Stroncek, D., Poston, L., Akel, S., Hankins, J.S., et al. (2022). Optimizing haematopoietic stem and progenitor cell apheresis collection from plerixafor-mobilized patients with sickle cell disease. Br. J. Haematol. 198, 740–744.

59. Bolger, A.M., Lohse, M., and Usadel, B. (2014). Trimmomatic: a flexible trimmer for Illumina sequence data. Bioinformatics 30, 2114–2120.

60. Magoč, T., and Salzberg, S.L. (2011). FLASH: fast length adjustment of short reads to improve genome assemblies. Bioinformatics 27, 2957–2963.

61. Li, H., and Durbin, R. (2009). Fast and accurate short read alignment with Burrows-Wheeler transform. Bioinformatics 25, 1754–1760.

## Supplemental References

1. Traxler, E. A. et al. A genome-editing strategy to treat β-hemoglobinopathies that recapitulates a mutation associated with a benign genetic condition. Nat. Med. 22, 987–990 (2016).

2. Métais, J.-Y. et al. Genome editing of HBG1 and HBG2 to induce fetal hemoglobin. Blood Adv 3, 3379–3392 (2019).

3. Giannoukos, G. et al. UDiTaS^TM^, a genome editing detection method for indels and genome rearrangements. BMC Genomics 19, 212 (2018).

